# Persistent Nav1.6 current drives spinal locomotor functions through nonlinear dynamics

**DOI:** 10.1101/2023.04.18.537411

**Authors:** Benoit Drouillas, Cécile Brocard, Sébastien Zanella, Rémi Bos, Frédéric Brocard

**Affiliations:** Institut de Neurosciences de la Timone, UMR 7289, Aix-Marseille Université and Centre National de la Recherche Scientifique (CNRS), Marseille, France; Lead Contact

**Keywords:** Posture, Locomotion, Spinal cord, Motoneuron, CPG, Bistability, *Nav1.6* channels.

## Abstract

Persistent sodium current (*I*_NaP_) in the spinal locomotor network promotes two distinct nonlinear firing patterns: a self-sustained spiking triggered by a brief excitation in bistable motoneurons and bursting oscillations in interneurons of the central pattern generator (CPG). Here, we identified the NaV channels responsible for *I*_NaP_ and their role in motor behaviors. We report the axonal *Nav1.6* as the main molecular player for *I*_NaP_ in lumbar motoneurons. The motoneuronal inhibition of *Nav1.6*, but not of *Nav1.1*, impairs *I*_NaP_, bistability, postural tone and locomotor performance. In interneurons of the CPG region, *Nav1.6* with *Nav1.1* equally mediate *I*_NaP_ and the inhibition of both channels is required to abolish oscillatory bursting activities and the locomotor rhythm. Overall, *Nav1.6* plays a significant role both in posture and locomotion by governing *I*_NaP_-dependent bistability in motoneurons and working in tandem with *Nav1.1* to provide *I*_NaP_-dependent rhythmogenic properties of the CPG.

## Introduction

The spinal central pattern generator (CPG) governs basic features of locomotor movements, with downstream motoneurons conveying CPG commands to muscles ^1, 2^. The rhythmogenic module of the CPG comprises excitatory interneurons endowed with inherent persistent Na^+^ current (*I*_NaP_)-mediated membrane oscillations at a frequency range similar to locomotor rhythms ^3–10^. The activity-dependent increase of *I*_NaP_ when the locomotor rhythm emerges ^5, 6^ or the impairment of the locomotor rhythm when *I*_NaP_ is inhibited in vertebrates ^8, 11–15^, support the fundamental role of *I*_NaP_ in driving the locomotor rhythm through the generation of neuronal oscillations in the CPG. Beyond its rhythmogenic role, *I*_NaP_ also amplifies synaptic transmission by enabling self-sustained spiking activity triggered in spinal motoneurons by a brief excitation ^16–18^. This bistable behavior has recently been shown to be a cellular correlate of the postural tone in hindlimbs ^19^. In sum, in agreement with motor dysfunctions often associated with its alteration ^20–26^, *I*_NaP_ appears to be critical for the operation of the spinal locomotor network in vertebrates.

Despite our progress in understanding the functional roles played by *I*_NaP_ within the spinal locomotor network, we still know little regarding the identity of Nav channel isoform(s) that give rise to *I*_NaP_. Here, we addressed this gap in knowledge by identifying *Nav1.6* as the molecular constituent of *I*_NaP_ in bistable motoneurons, and provided evidence of its critical role in promoting self-sustained spiking to support a postural tone in hindlimbs. We also found *Nav1.6* working in tandem with *Nav1.1* within the CPG to produce both *I*_NaP_-dependent oscillations at the neuronal level and the locomotor rhythm at the network level.

## Results

### *Nav1.6* expression predominates in lumbar motoneurons

To study the subcellular distribution of the two main sodium channels (*Nav1.1* and *Nav1.6*) of the ventral spinal cord ^27, 28^, we performed immunohistochemistry both in lumbar motoneurons (L_4_–L_5_) and interneurons of the locomotor CPG region (L_1_-L_2_). The pan-specific antibody (PanNav) highlighted all Nav channel isoforms along ankyrin-G positive axon initial segments (AIS; Figure S1A,B). The distribution of *Nav1.1* and *Nav1.6* channels is segregated in motoneurons (Figure 1A). The *Nav1.1* was expressed proximally within the first 10 µM of the PanNav staining while *Nav1.6* mostly paralleled the staining profile of the PanNav along the AIS (Figure 1A,B). Specifically, *Nav1.6* gradually increased in density in the first 20 µm and declined in the last 10 µm (Figure 1B). As a result, the PanNav colocalized more extensively with *Nav1.6* than with *Nav1.1* (Figure 1C). In interneurons of the locomotor CPG region located in ventromedial part of upper lumbar segments ^29, 30^, *Nav1.1* and *Nav1.6* were expressed mainly within the first and the second half of the PanNav staining, respectively (Figure 1D,E). The distribution of the PanNav staining uniform along the AIS results from this complementary gradient (Figure 1E). Thus, PanNav exhibited an equal colocalization with *Nav1.1* and *Nav1.6* (Figure 1F).

**Figure 1:**
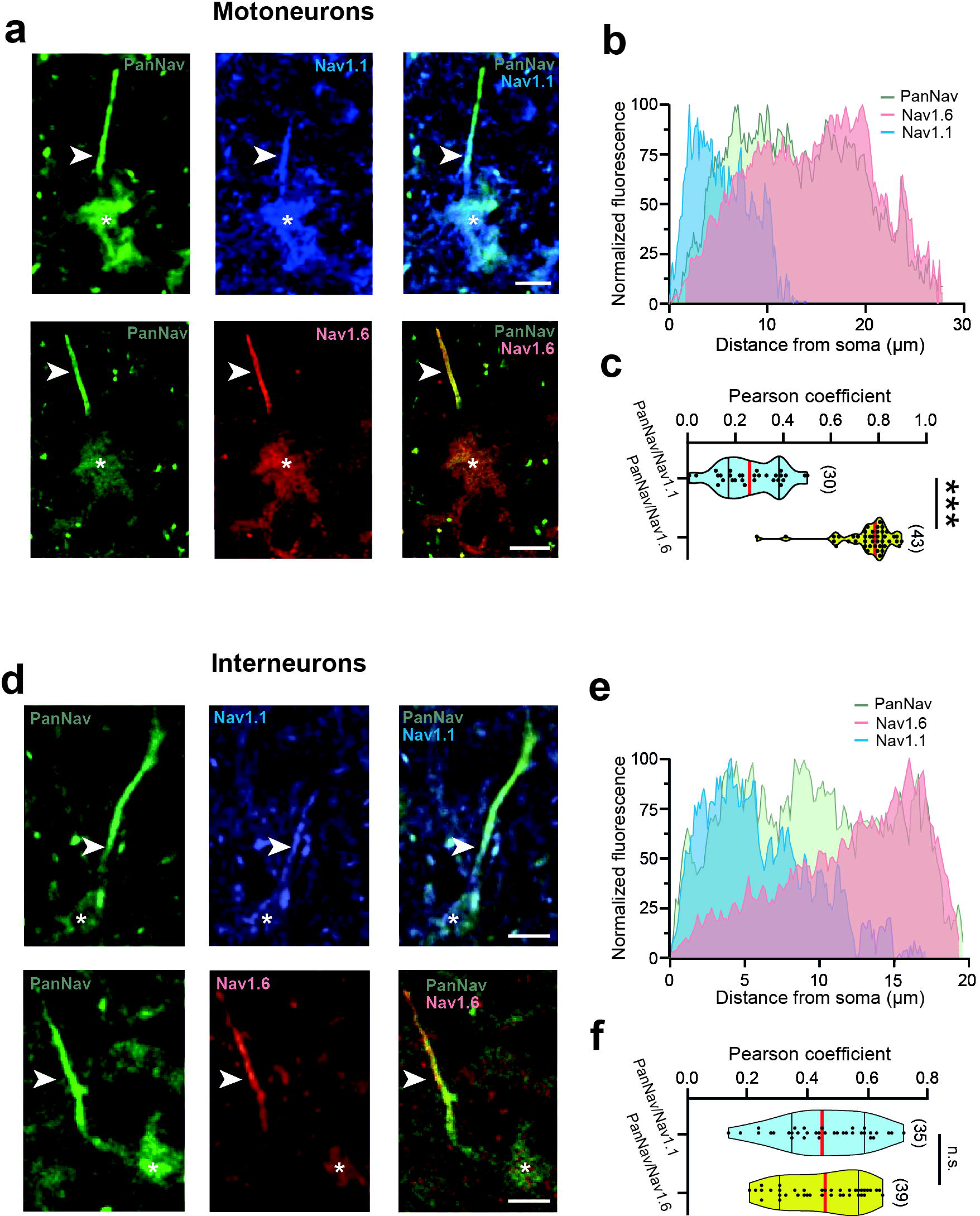
Distribution of *Nav1.1* and *Nav1.6* subunits in the axon initial segments (AISs) of lumbar motoneurons and interneurons of the locomotor CPG region. **a,d** Representative optical sections showing the immunostaining of all Nav α-subunits (left, green panels), *Nav1.1* subunits (middle-top, blue panels) or *Nav1.6* subunits (middle-low, red panels) along the AISs (arrowheads) of lumbar L4/L5 motoneurons (**a**) or L1/L2 ventromedial interneurons (**d**). The right panels are merged images. Scale bars represent 10 μm for **a** and 5 µm for **d**. **b,e** The mean fluorescence intensity profile for the PanNav (green), *Nav1.1* (blue) and *Nav1.6* (red) immunostainings along the PanNav-labelled AIS of motoneurons (**b**) and interneurons (**e**). For each antibody, the immunofluorescence was normalized to its maximum intensity. **c,f** Violin plots of the Pearson’s coefficient between PanNav and Na1.1 (cyan) or between PanNav and *Nav1.6* (yellow) in motoneurons (**c**) and interneurons (**f**). Numbers in brackets in **c,f** indicate the numbers of neurons. Each dot in **c** and **f** represents an individual neuron. n.s., no significance; \*\*\**P* < 0.001 (Unpaired t-test for **c,f**). For detailed *P* values see Source Data. Source data are provided as a Source Data file. See also Figures S1.

Altogether, the *Nav1.6* channels predominate in lumbar motoneurons, while the relative expression of *Nav1.1* and *Nav1.6* appears more balanced in interneurons of the CPG region.

### *Nav1.6* channels are instrumental for bistability in motoneurons

To explore the biological significance of *Nav1.1* and *Nav1.6* in firing properties of spinal neurons, we used ICA-121431 (ICA) and 4,9-anhydrotetrodotoxin (4,9-ahTTX) as blockers of *Nav1.1* and *Nav1.6*, respectively ^31–33^. We defined the specific and optimal concentration of ICA and 4,9-ahTTX as the highest dose that did not affect characteristics of the spike in *Nav1.1*^-/-^ and *Nav1.6*^-/-^ motoneurons, respectively. From a dose-response curve the optimal concentration was established at 350 nM for ICA and 200nM for 4,9-ahTTX (Figure S2). As we previously reported ^16, 19^, most of the large motoneurons (∼90 %) displayed a self-sustained spiking activity dependent on *I*_NaP_ and triggered by a brief excitation (Figure 2A). This bistability was insensitive to ICA but abolished by 4,9-ahTTX (Figure 2B-D). The *Nav1.6* blocker decreased the excitability of motoneurons (Figure 2E) and altered both waveform and dynamics of the spike more strongly than the *Nav1.1* blocker (Figure 2F). Specifically, the spike was smaller and displayed a higher threshold (Figure 2G,H). Furthermore, ICA and 4,9-ahTTX lowered the firing rate (Figure 2I-L). Notably, the *Nav1.6* blocker diminished the capacity to spike at high frequencies and led to a frank failure of the repetitive firing, not observed with ICA. The genetic deletion of *Nav1.1* or *Nav1.6* broadly recapitulated the distinct contributions of the two channels in firing properties of motoneurons, only one third of *Nav1.6*^-/-^ motoneurons remained bistable (Figure S3). In *Nav1.6*^-/-^ motoneurons, the expression of *Nav1.1* was not restrained to the proximal part of the AIS like in wildtype, but was spread along the full length of the AIS (Figure S1D,E). The *Nav1.1* colocalized more extensively with PanNav in *Nav1.6*^-/-^ than in wildtype motoneurons (Figure S1F). The compensation was independent of structural changes as the ankyrin-G expression pattern did not differ from wildtype (Figure S1C). A similar compensation by *Nav1.6* did not occur for *Nav1.1* in motoneurons (Figure S1G-I).

**Figure 2:**
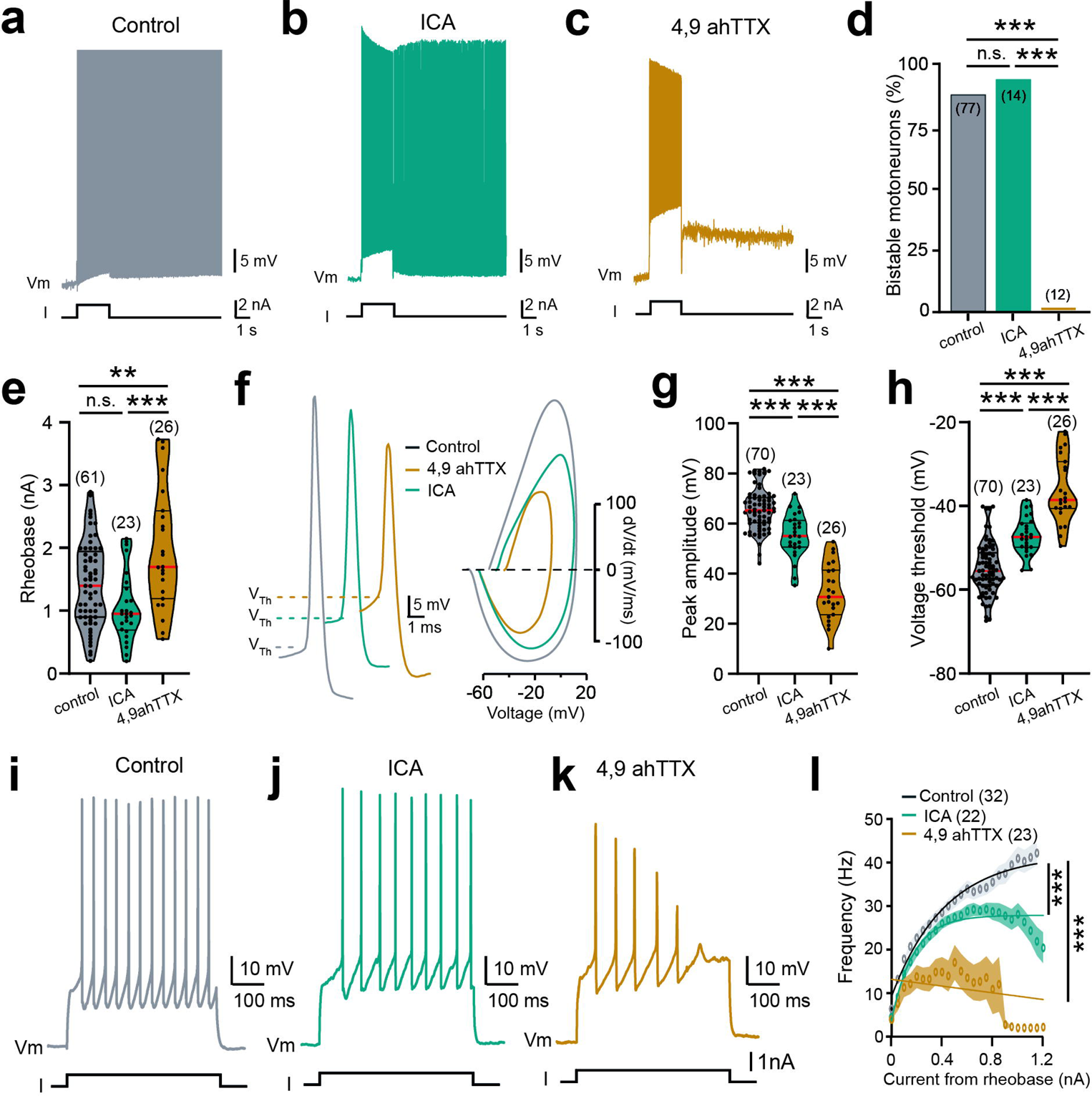
*Nav1.6* channels are instrumental for bistability in lumbar motoneurons. **a-c,f,i-k** Voltage traces from motoneurons recorded in response to suprathreshold (**a-c**), near-threshold (**f**) or incrementing (**i-k**) depolarizing pulses before (**a**,**f,i,** grey, *n* = 6 mice) or during bath application of ICA-121431 (**b,f,j**, green, ICA, 350 nM, *n* = 4 mice) or 4,9-anhydrotetrodotoxin (**c,f,k**, orange, 4,9 ahTTX, 200 nM, *n* = 4 mice). **d** Quantification of the proportion of bistable motoneurons. **e,g,h** Violin plots of the rheobase (**e**), peak amplitude (**g**) and threshold (**h**) of the action potential. **f** Representative individual action potentials (left) with their phase plots (right) generated from the first derivative (dV/dt; y-axis) versus membrane potential (mV; x-axis). Dashed lines indicate the spiking threshold (V_Th_). **l** Firing frequency as a function of the amplitude of the current pulse. Numbers in brackets in **d,e,g,h,l** indicate the number of motoneurons. Each dot in **e,g,h** represents an individual motoneuron. Continuous lines in **l** represent best fit functions for experimental data with 95% confidence interval. n.s., no significance; \*\**P* < 0.01; \*\*\**P* < 0.001 (two-tailed Fisher test for **d**; one-way ANOVA with multiple comparisons for **e,g,h**; comparison of the fits for **l**). For detailed *P* values, see Source data. Source data are provided as a Source data file. See also Figures S2 and S3.

Overall, these data indicate a critical role of *Nav1.6* channels in setting the excitability and firing properties of motoneurons, notably by supporting the self-sustained spiking activity that cannot be supported by *Nav1.1* channels.

### *Nav1.6* partnered with *Nav1.1* to generate bursting properties in interneurons

Most of interneurons (∼90 %) from the CPG region displayed nonlinear firing properties manifested by bursting activities dependent on *I*_NaP_ and triggered by removing the extracellular Ca^2+^ (Figure 3A,E) ^5, 6^. In nearly half of these interneurons, the pharmacological inhibition of either *Nav1.1* or *Nav1.6* abolished the bursting activity (Figure 3E). In the other half the robustness of bursts decreased both in amplitude and duration without affecting the frequency (Figure 3B,C,F-H). The simultaneous application of the two blockers was required to abolish all bursting cells (Figure 3D,E). Bursting activities in *Nav1.1*^-/-^ and *Nav1.6*^-/-^ mice remained robust, comparable to those recorded in wildtype mice, but were abolished by 4,9-ahTTX and ICA, respectively (Figure S4A-F). In wildtype interneurons recorded in normal aCSF, the pharmacological inhibition of one of the two Nav channels equally decreased the amplitude of the spike and depolarized the threshold (Figure 3I-K). Likewise, ICA and 4,9-ahTTX indistinguishably lowered the firing rate as the depolarizing current pulses increased (Figure 3L-O). Smaller spikes were also observed both in *Nav1.1*^-/-^ and *Nav1.6*^-/-^ interneurons (Figure S4G,H). However, the higher voltage threshold and the lower firing rate were observed only in *Nav1.1*^-/-^ interneurons (Figure S4I-M). Once again, partial compensatory mechanisms occurred in *Nav1.6*^-/-^ interneurons with the *Nav1.1* expression extended to the distal part of AIS (Figure S1J-L). Conversely, such homeostatic regulation by *Nav1.6* was not present in *Nav1.1*^-/-^ interneurons (Figure S1M-O).

**Figure 3:**
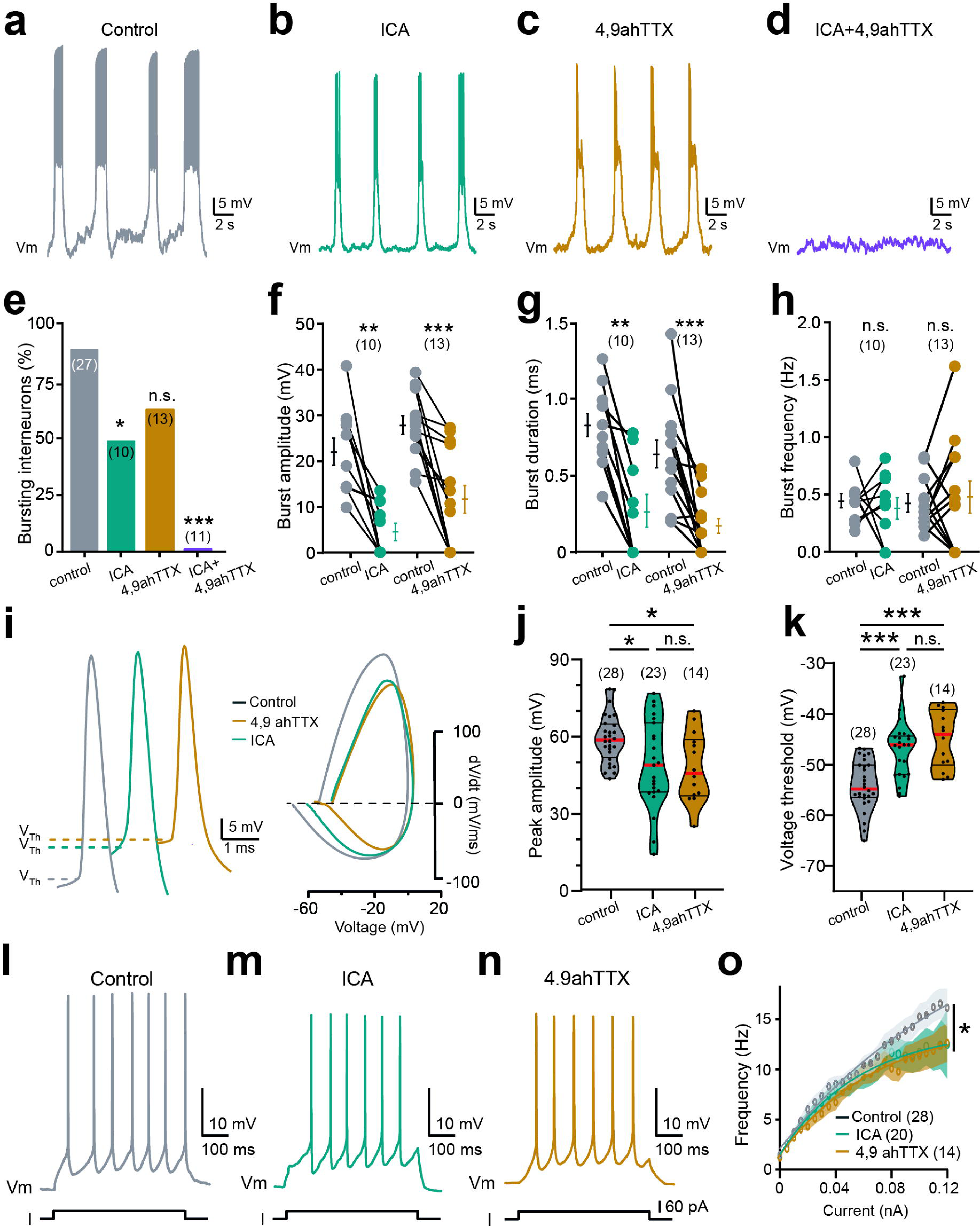
*Nav1.6* channels work in tandem with *Nav1.1* channels for generating *I*_NaP_-dependent bursting properties. **a-d** [Ca^2+^]_o_-free-saline-induced bursting activity recorded from ventromedial interneurons of the locomotor CPG region (L_1_-L_2_) before (**a,** *n* = 7 mice) or during the bath application of ICA-121431 (**b,** ICA, 350 nM, *n* = 5 mice) or 4,9-anhydrotetrodotoxin (**c,** 4,9 ahTTX, 200 nM, *n* = 6 mice) or when the two drugs were co-applied (**d,** *n* = 7 mice). **e** Quantification of the proportion of bursting cells. **f-h** Quantification of burst parameters. **i,l-n** Voltage traces recorded from interneurons of the locomotor CPG region in response to near-threshold (**i**) or incrementing (**l-n**) depolarizing pulses before (**i,l,** grey, *n* = 3 mice) or during the bath application of ICA-121431 (**i**,**m,** green, ICA, 350 nM, *n* = 3 mice) or 4,9-anhydrotetrodotoxin (**i,n,** orange, 4,9 ahTTX, 200 nM, *n* = 2 mice). **i** Representative individual action potentials (left) with their phase plots (right) generated from the first derivative (dV/dt; y-axis) versus membrane potential (mV; x-axis). Dashed lines indicate the spiking threshold (V_Th_). **o** Firing frequency as a function of the amplitude of the current pulse. Numbers in brackets in **e-h,j,k,o** indicate the numbers of interneurons. Each dot in **j,k** represents an individual interneuron. Continuous lines in **o** represent best fit functions for experimental data with 95% confidence interval. n.s., no significance; \**P* < 0.05; \*\**P* < 0.01; \*\*\**P* < 0.001 (two-tailed Fisher test for **e**; two-tailed Wilcoxon paired test for **f-h**; one-way ANOVA with multiple comparisons for **j,k**; comparison of the fits for **o**). Mean ± SEM. For detailed *P* values, see Source data. Source data are provided as a Source data file. See also Figure S4.

Together, these data show that both *Nav1.1* and *Nav.1.6* channels, ubiquitously expressed in the CPG region, are partners and take an equivalent part in regulating spiking properties and dynamics of oscillatory activities in interneurons.

### *I*_NaP_ dominantly relies on *Nav1.6* in motoneurons and on the cooperative gating between *Nav1.1* and *Nav1.6* in interneurons

We here examined the contribution of the two Nav channels in biophysical properties of *I*_NaP_. In response to a slow ramp voltage, motoneurons and interneurons typically displayed a large inward voltage-dependent current attributable to *I*_NaP_ (Figure 4 A,E, see also ^5^). In motoneurons, the *Nav1.1* blocker ICA did not significantly alter *I*_NaP_ (Figure 4A-D). By contrast, the *Nav1.6* blocker 4,9-anhydro-TTX strongly decreased *I*_NaP_ in amplitude and shifted toward more depolarized values its voltage activation threshold without modifying the half activation potential (Figure 4A-D). The co-application of the two drugs abolished *I*_NaP_ (Figure 4B). Data were recapitulated in *Nav1.1*^-/-^ and *Nav1.6*^-/-^ motoneurons (Figure S5A-D) and the residual *I*_NaP_ was abolished by 4,9-ahTTX and ICA, respectively (Figure S5E-H). In interneurons, in the presence of either ICA or 4,9-ahTTX, the peak amplitude of *I*_NaP_ was mostly halved without affecting its activation threshold or the half activation potential (Figure 4E-H). Likewise, the co-application of the two drugs abolished *I*_NaP_ (Figure 4F). Astonishingly, biophysical properties of *I*_NaP_ in *Nav1.1*^-/-^ and *Nav1.6*^-/-^ interneurons were similar to wildtype littermates (Figure S5I-L) but it was almost inhibited by 4,9-anhydro-TTX and ICA, respectively (Figure S5M-P).

**Figure 4:**
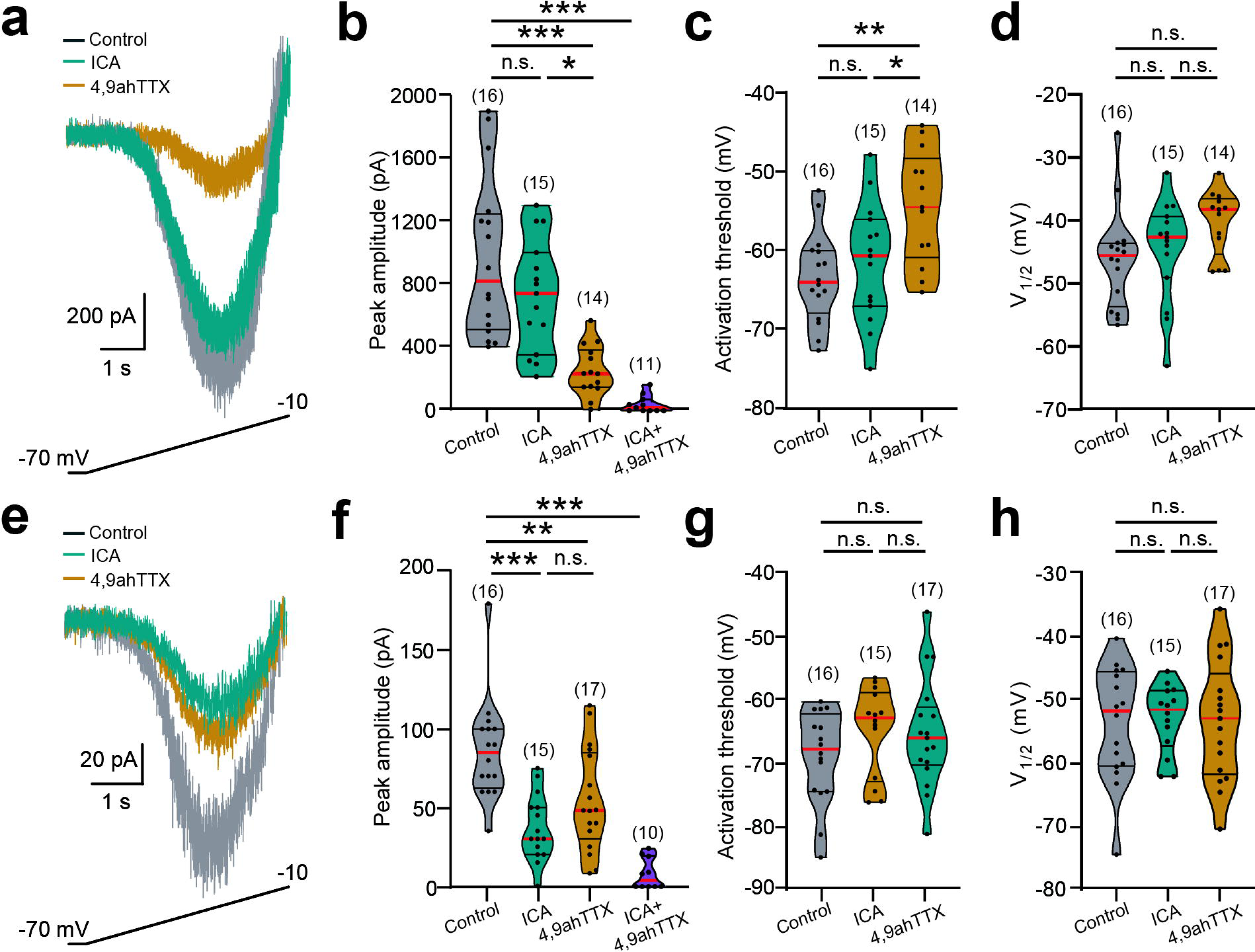
*Nav1.6* channels encode *I*_NaP_ in motoneurons. **a,e** Superimposed leak-subtracted *I*_NaP_ recorded from motoneurons (**a**) and interneurons of the CPG region (**e**) in response to a slow ramping depolarization before (grey, *n* = 6 mice) and after the bath application of ICA-121431 (green, ICA, 350 nM, *n* = 4 mice) or 4,9-anhydrotetrodotoxin (orange, 4,9 ahTTX, 200 nM, *n* = 4 mice). **b-d,f-h** Amplitude (**b,f**), threshold (**c,g**) and half-activation voltage (**d,h**) of *I*_NaP_. Numbers in brackets indicate the number of neurons. Each dot represents an individual neuron. n.s., no significance; \**P* < 0.05; \*\**P* < 0.01; \*\*\**P* < 0.001 (one-way ANOVA with multiple comparisons). For detailed *P* values, see Source data. Source data are provided as a Source data file. See also Figure S5.

These results further support the concept that *I*_NaP_ is dominantly mediated by *Nav1.6* in motoneurons and by the *Nav1.1*-*Nav1.6* partnership in interneurons.

### *Nav1.1* and *Nav1.6* channels participate in fictive locomotion

The presence of *I*_NaP_ at the level of the CPG is required for the locomotor rhythm generation in rodents ^3–6, 11, 12^. Since the *Nav1.1*-*Nav1.6* partnership generates *I*_NaP_ in the CPG region, we examined the role of the two channels in the operation of the locomotor rhythm-generating network by using whole-mount spinal cord preparation. During fictive locomotion, the bath-application of ICA or 4,9-ahTTX on rhythm-generating networks decreased the locomotor burst amplitude without affecting temporal parameters of locomotor outputs (Figure 5A-E). The co-application of the two drugs strongly impaired locomotor outputs and ultimately led to abolish them in most preparations (Figure 5F-H). Note that the spinal cords isolated from *Nav1.1*^-/-^ or *Nav1.6*^-/-^ mice exhibited a quite normal locomotor-like activity (Figure S6A,B).

**Figure 5:**
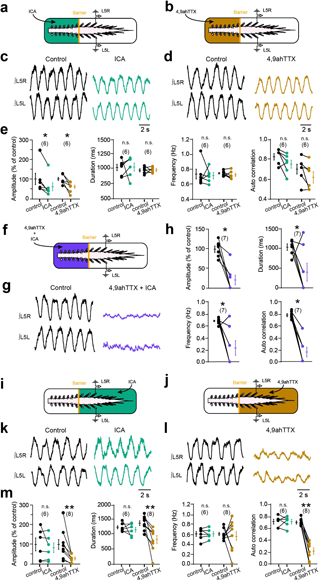
*Nav1.1* and *Nav1.6* channels participate in fictive locomotion. **a,b,f,i,j** Schematic representation of the whole-mount spinal cord with the recording glass electrodes from the lumbar segment L_5_ (L_5_R/L). The yellow solid line represents the Vaseline barrier. Built at L_2_/L_3_ level the Vaseline barrier allows the selective application of drugs over the rostral (**a,b,f,** above the L_3_ segment) or the caudal lumbar segments (**i,j,** below L_3_ segment). **c,d,g,k,l** Ventral-root recordings of NMA/5-HT-induced fictive locomotor activity before and after adding ICA-121431 (**c,k,** ICA, 350 nM, *n* = 12 mice) or 4,9-anhydrotetrodotoxin (**d,l,** 4,9 ahTTX, 200 nM, *n* = 14 mice) or the two drugs (**g,** *n* = 7 mice) to rostral (**c,d,g**) or caudal (**k,l**) lumbar segments. **e,h,m** Quantification of locomotor burst parameters. Numbers in brackets in **e,h,m** indicate the number of spinal cords. n.s., no significance; \**P* < 0.05; \*\**P* < 0.01 (two-tailed Wilcoxon paired test for **e,h,m**). Mean ± SEM. For detailed *P* values see Source Data. Source data are provided as a Source Data file. See also Figure S6.

The recurrent recruitment of *I*_NaP_ in motoneurons has also been shown to increase the amplitude of the locomotor drive from the CPG ^11^. To test the role of motoneuron *Nav1.1* and *Nav1.6* channels in the integration of rhythmic locomotor inputs we applied drugs to motoneuron pools caudally located (Figure 5I,J). The *Nav1.1* blocker ICA did not affect fictive locomotor outputs (Figure 5K,M). By contrast, the *Nav1.6* blocker 4,9-ahTTX strongly decreased the amplitude, duration and regularity of locomotor outputs without affecting the cycle period (Figure 5L,M). Note that these effects were observed without noticeable alterations of locomotor outputs recorded at the CPG level spared from blockers by the Vaseline barrier (Figure S6C-F).

Together, these data indicate that *Nav1.1* and *Nav1.6* taken individually are not essential for the locomotor rhythm generation, but they work in tandem at the level of the CPG to ensure rhythmogenesis. Our findings also suggest that *Nav1.6* channels are rhythmically activated in motoneurons to amplify locomotor outputs.

### Silencing *Nav1.6* in spinal motoneurons disturbs motor behaviors

To specifically discriminate the behavioral role of *Nav1.6* in spinal motoneurons, we injected the AAV2-retro encoding the *Nav1.6*-shRNA with a green fluorescent reporter (eGFP) into the *triceps surae* muscle of neonatal mice. The AAV2-retro was previously shown to high-efficiently, extensively and bilaterally transduce most of lumbar motoneurons after the unilateral intramuscular injection in neonatal mice ^34^. As early as the 2^nd^ week post-injection, the expression of eGFP was observed in ventral horns of the spinal cord from T7-T11 to S1-S3 and 46.8 ± 8.8 % of lumbar motoneurons (533 out of 1169 large cholinergic neurons in the L3-L5 ventral horns from 3 mice), were transduced (Figure 6A). Concomitantly, the viral transfection led to a decrease of the membrane protein expression of sodium channels in the lumbar spinal cord by ∼ 28 % (Figure 6B). The weak fixation condition required for immunostaining *Nav1.6* channels is not sufficient to retain the cytosolic GFP reporter from transduced cells. Despite this constraint we observed that one third (35 %) of lumbar motoneurons no longer expressed *Nav1.6* at the AIS (Figure 6C-F). Compared to motoneurons transduced with the scramble shRNA, the proportion of bistable motoneurons markedly decreased to ∼ 23 % in *Nav1.6-*shRNA mice (Figure 7A-C) with a significant decrease of *I*_NaP_ amplitude without modifying the activation threshold (Figure 7D-F). The amplitude of the spike and firing rate also decreased without affecting the spike threshold or the excitability of motoneurons (Figure 7G-M).

**Figure 6:**
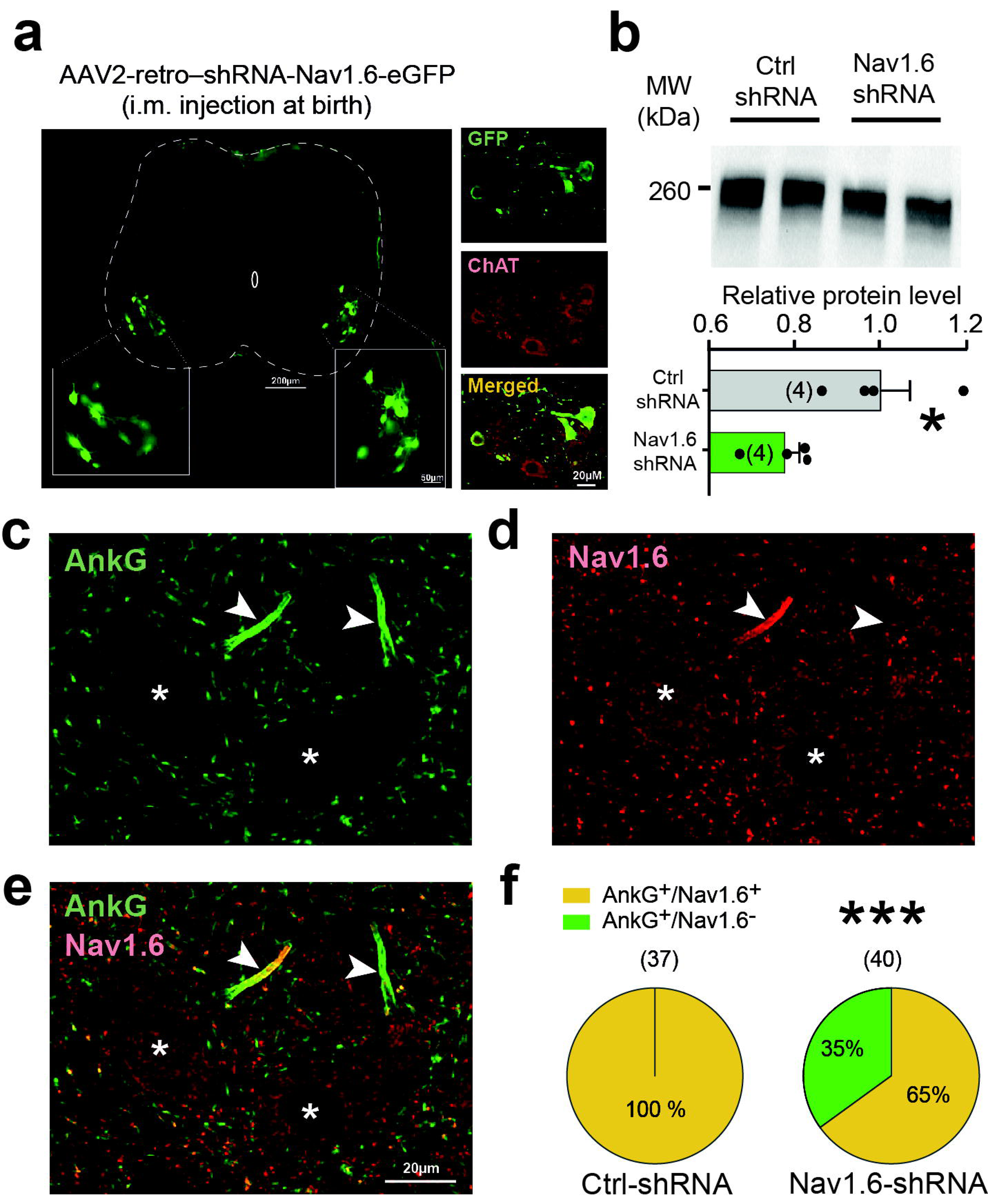
Silencing *Nav1.6* channels in spinal motoneurons. **a** Left: Native GFP-expressing motoneurons visualized in the lumbar spinal cord from a P10 mice and transduced by the AAV2-retro-shRNA-*Nav1.6*-eGFP injected into the *triceps surae* muscle at birth. Scale bar = 200 μm. Insets are high magnifications of the ventral horn showing native GFP-expressing motoneurons. Scale bar represents 50 μm. Right: High magnification of the ventral horn showing native fluorescence of motoneurons transduced by AAV2-retro (upper) and immunostained for choline acetyltransferase (middle, ChAT antibody; bottom, merged images). **b** Up: PanNav immunoblots of lumbar segments from P10 mice intramuscularly injected at birth with the AAV2-retro encoding either a scramble shRNA (*n* = 4 mice) or a *Nav1.6*-targeting shRNA (*n* = 4 mice). One mouse per lane. Bottom: group means quantification of the ∼ 260 kDa band normalized to scramble-injected controls. **c-e** Representative single optical sections showing immunostaining against ankyrin-G (**c**) and *Nav1.6* (**d**) expressed in AISs of motoneurons from a P14 mice transduced by the *Nav1.6*-targeting shRNA; merge image in **e**. Asterisks indicate the motoneuron nucleus position and arrows AISs. Scale bars represent 20 µm. **f** Proportion of motoneurons double-labelled with ankyrin-G and *Nav1.6* antibodies in mice transduced with the scramble shRNA (left, *n* = 3 mice) or with the *Nav1.6*-shRNA (right, *n* = 3 mice). Numbers in brackets in **f** indicate the numbers of motoneurons. \**P* < 0.05; \*\*\**P* < 0.001 (two-tailed Mann-Whitney test for **b**; two-tailed Fisher test for **f**). Mean ± SEM. For detailed *P* values see Source Data. Source data are provided as a Source Data file.

**Figure 7:**
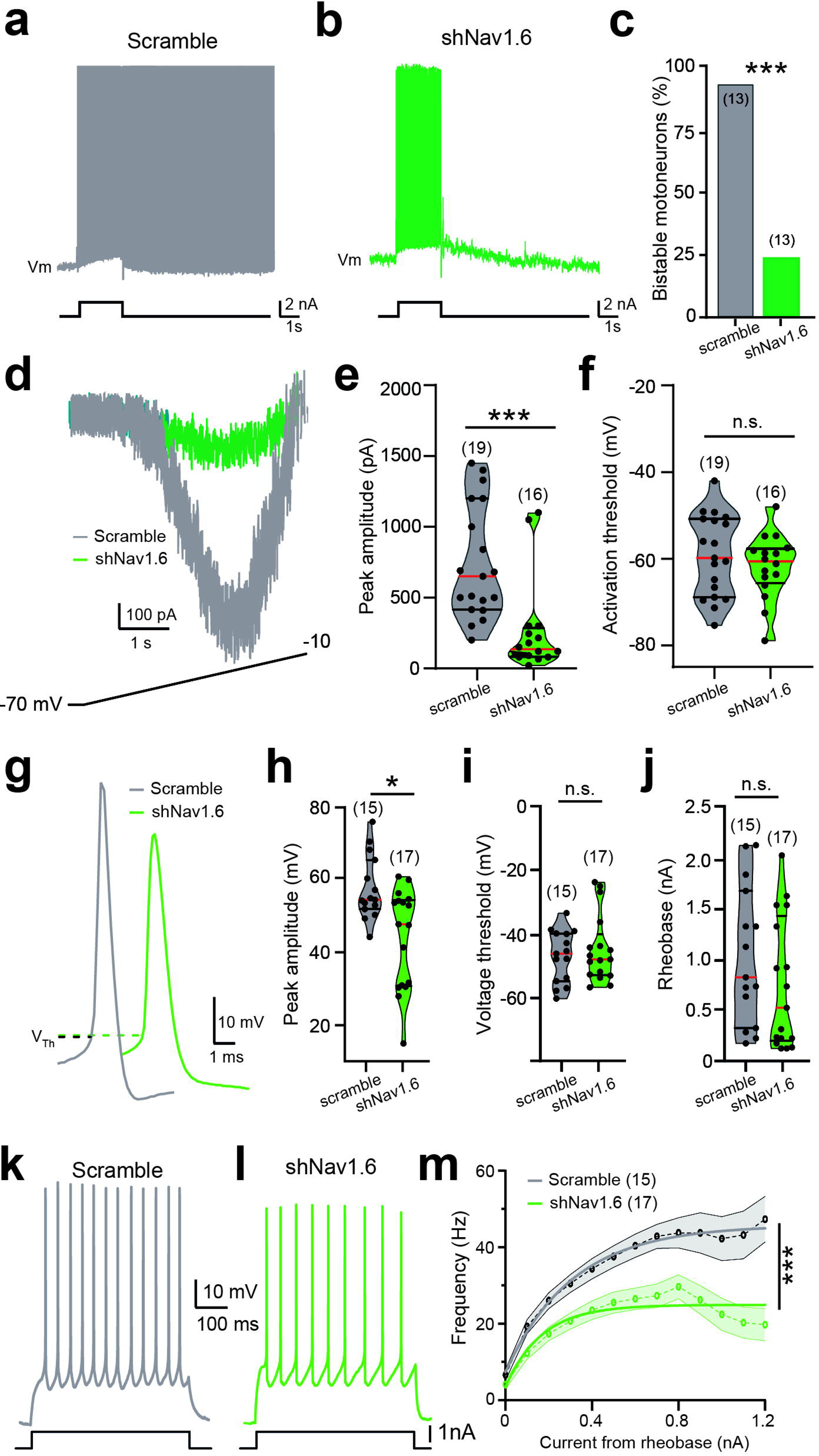
Silencing *Nav1.6* impairs the self-sustained spiking activity of lumbar motoneurons. **a,b,g,k,l** Voltage traces in response to suprathreshold (**a,b**), near-threshold (**g**) or incrementing (**k,l**) depolarizing pulse and recorded from eGFP+ motoneurons transduced either with the scramble (**a,g**,**k** grey, *n* = 3 mice) or with the *Nav1.6*-targeting shRNA (**b,g**,**l** green, *n* = 4 mice). **c** Group mean quantification of the proportion of bistable motoneurons. **d** Superimposed leak-subtracted *I*_NaP_ recorded in response to a slow ramping depolarization from eGFP+ motoneurons transduced either with the scramble (grey, *n* = 2 mice) or with the *Nav1.6*-targeting shRNA (green, *n* = 2 mice). **e,f** Violin plots of the peak amplitude (**e**) and voltage threshold (**f**) of *I*_NaP_. **h,i,j**, Violin plots of the peak amplitude (**h**) voltage threshold (**i**) and rheobase (**j**) of the action potential. **m** Firing frequency as a function of the amplitude of the current pulse. Continuous lines represent best fit functions for experimental data with 95% confidence interval. Numbers in brackets in **c,e,f,h-j,m** indicate the number of recorded motoneurons. Each circle represents an individual motoneuron. n.s., no significance; \**P* < 0.05; \*\*\**P* < 0.001 (two-tailed Fisher test for **c**; two-tailed Mann-Whitney test for **e,f,h-j**; comparison of the fits for **m**). For detailed *P* values see Source Data. Source data are provided as a Source Data file.

Mice transduced with *Nav1.6*-shRNA can be phenotypically distinguished from control-shRNA mice as early as the 2^nd^ week post-injection. At rest, *Nav1.6*-shRNA mice typically showed a wider base of support with hind paws pointed outward and visible outside the body contour (Figure 8A). Besides having reduced hind paw print areas (Figure 8B), *Nav1.6*-shRNA mice showed striking locomotor deficits as they failed to adapt to accelerated speed in the rotarod test (Figure 8C). These locomotor deficits did not fade over weeks and persisted in young (4 wk) adult mice (Figure 8C). At this age, our data indicate that temporal parameters of the stepping movements were normal in *Nav1.6*-shRNA mice, with animals walking with a regular and alternating locomotor pattern, although with a wider base of support (Figure 8D). However, *Nav1.6*-shRNA mice stereotypically walked with a markedly collapsed basin (Figure 8E and Videos S1 and S2). Kinematic reconstruction of hindlimb movements revealed that hip and knee joint angles from *Nav1.6*-shRNA mice were ∼ 10° less extended throughout the stance phase, while ankle joint angles remained similar to control-shRNA mice (Figure 8F,G). No significant differences were observed in kinematic variables during the swing phase (Figure 8F,G).

**Figure 8:**
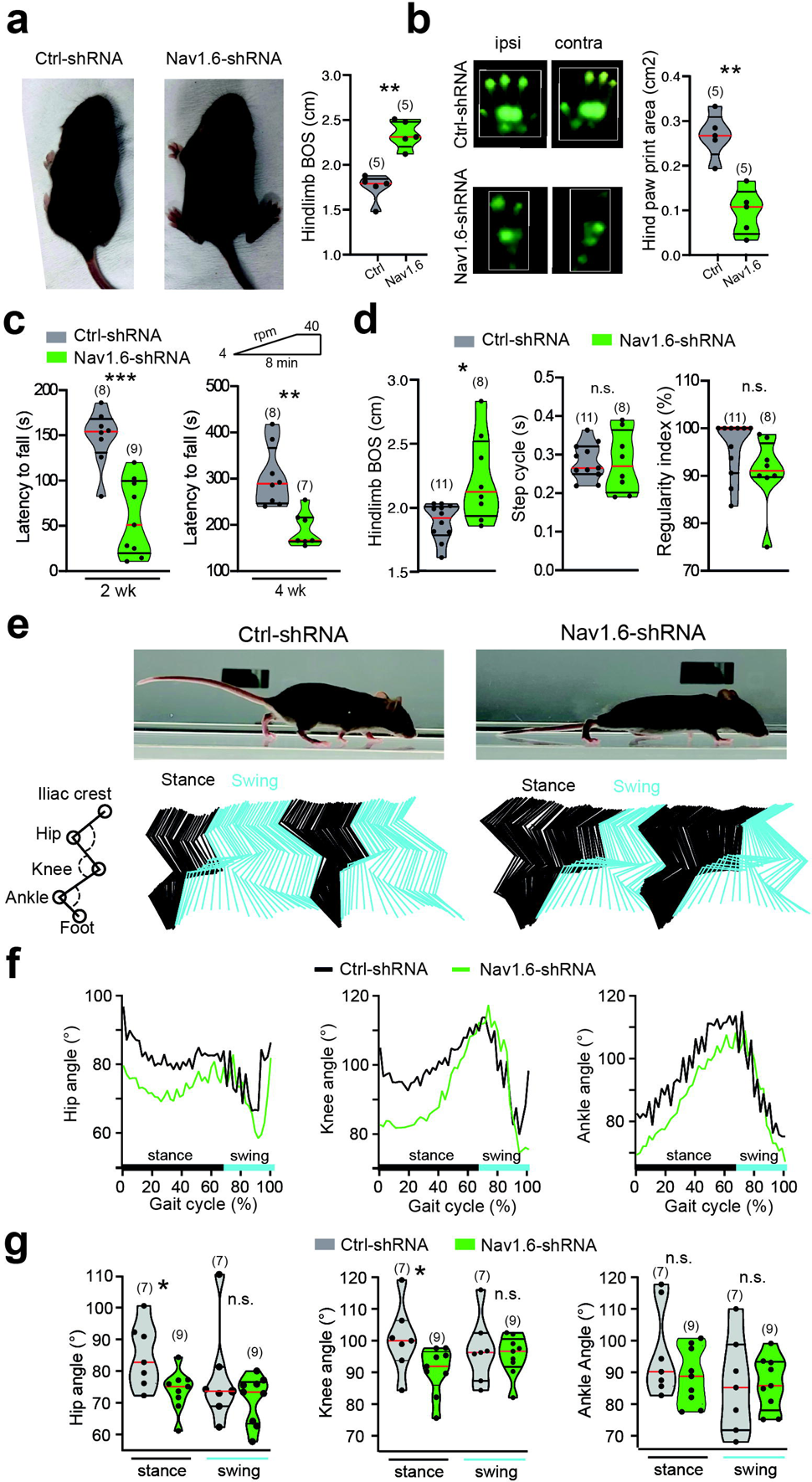
Silencing *Nav1.6* in lumbar motoneurons disturbs motor behaviors. **a** left and middle: Top views of 2 wk-old wildtype mice transduced either with the scramble shRNA (left) or with the *Nav1.6*-shRNA (middle). Right: Violin plots of the base of support at rest. **b** left: Typical hind paw prints captured by the CatWalk XT automated gait analysis system from 2-wk-old wildtype mice either transduced with the scramble shRNA (top) or with the *Nav1.6*-shRNA (bottom). Right: Violin plots of the hind paw print area. **c** Violin plots of the latency to fall from a rod rotating at accelerated speed (4 to 40 rpm). The mice were stratified into two age-groups: 2-and 4-week-old mice (2wk, 4wk). **d** Violin plots of the base of support (left), step cycle (middle) and regularity index of paw placements (right) during CatWalk locomotion from 4-week-old mice. **e** Side view of 4 wk-old wildtype mice freely walking through a corridor on a glass walkway and transduced either with the scramble shRNA (left) or with the *Nav1.6*-shRNA (right). bottom: Color-coded stick decomposition of hindlimb movement during two consecutive steps. **f** angular excursion of three joints (hip, knee and ankle) during the locomotor cycle. **g** Decoding performance for three joint angles {mean θ of the hip, knee and ankle} during the stance and swing phases. Numbers in brackets indicate the number of mice. Each circle represents an individual mouse. n.s., no significance; \**P* < 0.05; \*\**P* < 0.01; \*\*\**P* < 0.001 (two-tailed Mann–Whitney test for **a,b,c,d,g**). For detailed *P* values, see Source data. Source data are provided as a Source data file. See also Videos S1 and S2.

Altogether, these data provide evidence that *Nav1.6*, as the molecular constituent of *I*_NaP_ in lumbar motoneurons, is critical in promoting self-sustained spiking activity and plays a key role in producing an efficient postural tone in hindlimbs.

## Discussion

The study provides important insights into how the spinal locomotor network operates: i) it shows that *Nav1.6* works in tandem with *Nav1.1* in CPG interneurons to generate *I*_NaP_-bursting and thereby the locomotor rhythm, ii) it identifies *Nav1.6* as the main molecular player for *I*_NaP_ in bistable motoneurons to promote a self-sustained spiking activity, iii) it assigns to motoneuronal *Nav1.6* channels behavioral roles in producing a postural tone and amplifying the locomotor output.

Out of the nine α-subunits, *Nav1.1*, *Nav1.2, Nav1.3* and *Nav1.6* predominate in the CNS ^35^. We focused on *Nav1.1* and *Nav1.6* because of their strong expression within the ventral spinal cord ^27, 28, 36–38^, while *Nav1.2* and *Nav1.3* are predominantly found in dorsal horn sensory neurons ^27, 28, 38^. We identified *Nav1.1* and *Nav1.6* as the molecular determinants for most of the *I*_NaP_ within the spinal locomotor network. If *Nav1.1* and *Nav1.6* conduct *I*_NaP_ in the CPG region, *Nav1.6* makes a larger contribution to *I*_NaP_ in motoneurons. This agrees with the predominant expression of *Nav1.6* in the AIS of motoneurons, supposed to be the primary source of *I*_NaP_ ^39^. As also found in supraspinal structures ^40–44^, *Nav1.6* controls the excitability of spinal neurons by hyperpolarizing the spike threshold. The *I*_NaP_ hyperpolarizes the spike threshold both in motoneurons and CPG interneurons ^12^. Since *Nav1.6* singularly mediates *I*_NaP_ ^42, 45–55^, it might be assumed that the persistent *Nav1.6* current regulates neuronal excitability by lowering the spike threshold, especially in motoneurons ^56^.

In vertebrates, *I*_NaP_ functions as the primary mechanism to produce the locomotor rhythm ^4^ and we here show that *Nav1.1* and *Nav1.6* cooperate to mediate *I*_NaP_ within the CPG region. Both are necessary and sufficient for rhythmogenesis but the combination of the two is required for a robust locomotor rhythm. When applied over the caudal-most lumbar segments, the *Nav1.6* blocker 4,9-ahTTX reduces locomotor outputs similarly to the *I*_NaP_ blocker riluzole ^11, 12^. It is thus believed that the persistent *Nav1.6* current in motoneurons potentiates synaptic drives from the CPG to amplify locomotor outputs. This is highly plausible considering the role of I_NaP_ membrane oscillation amplification in motoneurons ^7^ and in initiating spikes during locomotor-like inputs ^57^. As fast walking is more difficult without *Nav1.6* in motoneurons, such mechanisms appear critical for high-demanding locomotor tasks. It is conceivable that the recruitment of *Nav1.6* channels in motoneurons rises with the speed of locomotion, amplifying drives from the CPG to produce forceful motor outputs.

The acute pharmacological inhibition of Nav channels differs qualitatively from the chronic deletion in genetic models. Specifically, the striking robustness of both bursting cells and the locomotor rhythm in mutant mice suggests long-term compensatory mechanisms. A shift from Na^+^-to Ca^2+^-dependent mechanism of bursting reported in *Nav1.6*^-/-^ Purkinje neurons ^58^ is unlikely in our Ca^2+^-free recording solution. A reciprocal homeostatic expression between Nav channel isoforms may contribute ^50, 51, 59, 60^ but does not appear as the main factor in our study. An alternative mechanism might involve a lower recruitment of K^+^ currents in response to the decrease of the Na^+^ conductance ^41, 58^. This hypothesis appeas to be the most likely since rhythmogenesis of the CPG relies on dynamic interactions between *I*_NaP_ and the M-current mediated by *Kv7.2* channels ^9^. Because Kv7.2 channels are co-expressed with Nav channels in CPG interneurons ^9^, we posit that *Nav1.1* and *Nav1.6* are key partners of Kv7.2 in regulating responsiveness of locomotor-related bursting cells.

Our finding shows *Nav1.6* as a key player of the bistability in motoneurons, while *Nav1.1* channels have a secondary role. The reduction in self-sustained spiking activity may be due to the unique biophysical properties of *Nav1.6* to exhibit a use-dependent potentiation during a long-lasting discharge ^61^, contrary to *Nav1.1* that inactivates ^62^. Once again, we posit the persistent *Nav1.6* current component as critical. This assumption follows earlier studies where bistable motoneurons disappear with the *I*_NaP_ blocker riluzole ^16, 17, 20^, irrespectively of the spike threshold shift induced by riluzole ^12, 57, 63, 64^. Therefore, the substantial drop in *I*_NaP_ following the deletion of *Nav1.6* offers a biophysical basis for the lower capacity of motoneurons to spike tonically. Collectively, a scenario arises where the persistent *Nav1.6* current leads to a mechanism of self-maintaining motoneuronal output as follow: secondary to the recruitment of Ca^2+^ channels during repetitive spikes, *Nav1.6*-mediated *I*_NaP_ indirectly fuel motoneurons in Ca^2+^ to activate Trpm5 channels governing a plateau potential responsible for the self-sustained spiking ^16, 19^. By virtue of this recruitment cascade, *Nav1.6* channels serve as the gateway to self-sustained spiking allowing efficient postural tone in muscles.

*Nav1.6* is expressed throughout the nervous system but not in muscles ^28, 36, 65, 66^. Its global deletion develops ataxia and tremors of hindlimbs progressing to paralysis ^67^. The disrupted function of the motor system likely involves central motor control impairments, but not of the CPG for locomotion since it operates normally in *Nav1.6*^-/-^ mice (Figure S6A,B). The selective inactivation of *Nav1.6* in cerebellar Purkinje neurons results in ataxia and tremor symptoms without paresis of hindlimbs ^43^. Here, the selective loss of *Nav1.6* in a large proportion of lumbar motoneurons decreases the hindlimb postural tone without ataxia or tremors. Then, the neurological disorders of *Nav1.6*^-/-^ mice can be decoupled and respectively attributed to distinct alterations in cerebellar (ataxia, tremor) and spinal (paresis of hindlimbs) neurons. We recently related the postural tone deficit to a decrease in the ability of motoneurons to produce a self-sustained spiking ^19^. From the present study, a straightforward conclusion is that *Nav1.6* supports most of the native *I*_NaP_ involved in promoting self-sustained spiking activity in bistable motoneurons. Therefore, the postural deficit after deleting or silencing *Nav1.6* channels can be considered a direct consequence of the loss of bistable motoneurons. According to a cell type-dependent expression of sodium channels ^68^, an heterogeneous role of *Nav1.6* channels throughout the motoneuronal population might explain the proximo-distal gradient motor deficit observed in *Nav1.6*-shRNA mice, reminiscent of the proximal muscle weakness reported in *Nav1.6*^-/-^ mice ^69^.

Overall, this study provides new insights into the operation of the locomotor network whereby *Nav1.1* and *Nav1.6* represent a functional set of voltage-gated sodium channels that endow the locomotor CPG with rhythmogenic properties, whereas a powerful motor output during locomotion is associated with the cyclic activation of *Nav1.6* channels in motoneurons. It also brings a clear support to a specific behavioral role of *Nav1.6* in motoneurons in controlling the postural tone by providing *I*_NaP_-dependent bistable properties.

## Author Contributions

B.D. designed, performed analyzed most of *in vitro* and *in vivo* experiments and wrote the first draft of the manuscript. C.B. designed, performed and analyzed immunohistochemistry and biochemistry experiments. S.Z. and R.B. designed and performed some of the *in vitro* experiments. F.B. conceptualized, administrated, designed, supervised and funded the whole project, performed and analyzed some *in vitro* experiments and wrote the manuscript.

## Acknowledgments

We are grateful to Jérémy Verneuil for coding videos, Hélène Bras for technical advices in the administration of the viral vector, Geneviève Rougon, Nejada Dingu for their valuable input to the manuscript. This research was supported by Agence National de la Recherche Scientifique (SpasT-SCI-T, ANR-21-CE17-0060 to F.B.), the French Institut pour la Recherche sur la Moelle épinière et l’Encéphale (to F.B.).

## Inclusion and diversity

We support inclusive, diverse and equitable conduct of research.

## STAR Methods

### Resource availability

#### Lead contact

Further information and requests for resources and reagents should be directed to and will be fulfilled by the Lead Contact, Frédéric Brocard.

#### Materials availability

This study did not generate new unique reagents.

#### Data and code availability

The published article includes all data generated or analyzed during this study.

### Experimental model and subject details

#### Mice

Mice (C57/Bl6 background) of either sex (P2-P3 for fictive locomotion experiments, P8-P14 for patch-clamp recordings, P14-P28 for behavioral experiments) were housed under a 12h light/dark cycle with *ad libitum* access to water and food. Room temperature was kept between 21-24°C and between 40-60% relative humidity. Mice heterozygous for the Scn1a (Nav1.1^+/-^) and Scn8a^med^ (*Nav1.6*^+/-^) were purchased from the Jackson laboratory (Bar Harbor, ME). Only homozygous animals were studied. All animal care and use were in compliance with the French regulations (Décret 2010-118) and approved by the ethics committee (Comité d’Ethique en experimentation animale, CEEA-071 Nb A1301404, authorization Nb 2018110819197361).

### Method details

#### shRNA construct

Specific shRNA sequence designed to knockdown *Nav1.6* transcript (TTGTCCTGAACACACTATTTA) was incorporated into a recombinant adeno-associated viral vector serotype 2 retrograde (rAAV2-retro), which features a U6 polymerase promoter to drive shRNA expression and a CMV promoter to drive eGFP expression for identification of transduced neurons (Vector Builder, Chicago, IL). We also used a non-targeting shRNA sequence (CCTAAGGTTAAGTCGCCCTCG) which has no homology to any known genes in mouse as a control. The standard titers of AAVs were ≥1 x 10^13^ GC/ml (genome copies/ ml).

#### Intramuscular vector delivery

The rAAV2-retro has been reported to extensively and high-efficiently transduce spinal motoneurons following a single intramuscular injection in neonatal mice ^34^. This technique was therefore chosen by unilaterally injecting the *triceps surae* muscle. Briefly, in pups cryoanesthetized at P2, the tip of the microcapillary preloaded with the AAV particles was lowered into the center of the muscle under optical microscope control. A total volume of 1.5 µL /animal was then slowly injected by hand with the micropipette held in place for an additional 30 s after injection before being slowly retracted.

#### *In vitro* preparations

*<u>For the slice preparation</u>*, the lumbar spinal cord was isolated in ice-cold (+4°C) artificial CSF (aCSF) solution composed of the following (in mM): 252 sucrose, 3 KCl, 1.25 NaH_2_PO_4_, 4 MgSO_4_, 0.2 CaCl_2_, 25 NaHCO_3_, 20 D-glucose, pH 7.4. The lumbar spinal cord was then introduced into a 1% agar solution, quickly cooled, mounted in a vibrating microtome (Leica, VT1000S) and sliced (325 µm) through the L1-L2 or L3-L5 lumbar segments. Slices were immediately transferred into the holding chamber filled with bubbled (95% O_2_ and 5% CO_2_) aCSF solution composed of (in mM): 120 NaCl, 3 KCl, 1.25 NaH_2_PO_4_, 1.3 MgSO_4_, 1.2 CaCl_2_, 25 NaHCO_3_, 20 D-glucose, pH 7.4, 30-32°C. After a 30-60 min resting period, individual slices were transferred to a recording chamber continuously perfused with aCSF heated to 32-34°C. *<u>For the whole-spinal cord preparation</u>,* the spinal cord was transected at T8-9, isolated and transferred with intact dorsal and ventral roots to the recording chamber. The tissue was continuously bubbled (95% O_2_ and 5% CO_2_) and perfused with heated (∼27-28°C) aCSF solution composed of (in mM): 120 NaCl, 4 KCl, 1.25 NaH_2_PO_4_, 1.3 MgSO_4_, 1.2 CaCl_2_, 25 NaHCO_3_, 20 D-glucose, pH 7.4.

#### In vitro recordings

*<u>For the slice preparation</u>*, whole-cell patch-clamp recordings were performed using a Multiclamp 700B amplifier (Molecular Devices) from L_3_-L_5_ motoneurons with the largest soma (>400µm^2^) located in the lateral ventral horn or from L_1_-L_2_ ventromedial interneurons adjacent to the central canal, a region proposed to contain a large part of the rhythm-generating locomotor network ^30^. Patch electrodes (2-4 MΩ) were pulled from borosilicate glass capillaries (1.5 mm OD, 1.12 mm ID; World Precision Instruments) on a Sutter P-97 puller (Sutter Instruments Company) and filled with intracellular solution containing (in mM): 140 K^+^-gluconate, 5 NaCl, 2 MgCl_2_, 10 HEPES, 0.5 EGTA, 2 ATP, 0.4 GTP, pH 7.3 (280 to 290 mOsm).

Pipette and neuronal capacitive currents were canceled and, after breakthrough, the series resistance was compensated and monitored. Recordings were digitized on-line and filtered at 10 kHz through a Digidata 1322A interface using Clampex 10.3 software (Molecular Devices). The main characterization of I_NaP_ was accomplished by slow ramp increase from −70 mV to −10 mV, slow enough (12 mV/s) to prevent transient sodium channel opening. All experiments were designed to gather data within a stable period (i.e., at least 5 min after establishing whole-cell access). *<u>For the whole spinal cord preparation</u>*, motor outputs were recorded from lumbar ventral roots by means of glass suction electrodes connected to an AC-coupled amplifier. The ventral root recordings were amplified (×1,000), high-pass filtered at 100 Hz, low-pass filtered at 5 kHz, and sampled at 10 kHz. Custom-built amplifiers enabled simultaneous online rectification and integration (100 ms time constant) of raw signals. Locomotor-like activity was induced by a bath application of N-methyl-DL aspartate (NMA, 10 μM) and 5-hydroxytryptamine (5-HT, 10 μM). In some experiments, a Vaseline barrier was built at the L_2_/L_3_ level to superfuse the highly rhythmogenic region of the locomotor network located in the rostral lumbar cord independently from the lumbar enlargement where most of motoneurons are present.

#### Assessment of motor behaviors

*<u>Walking.</u>* The CatWalkXT (Noldus Information Technology, Netherlands) was used to measure walking performance. Each animal walked freely through a corridor on a glass walkway illuminated with beams of light from below. A successful walking trial was defined as having the animal walk at a steady speed (no stopping, rearing, or grooming), and three to five successful trials were collected per animal. Footprints were recorded using a camera positioned below the walkway while a second camera placed parallel to the corridor captured hindlimb movements of the right side. The paw print and kinematic parameters were then analyzed using the CatWalk and the deepLabCut softwares, respectively. *<u>Rotarod test.</u>* Mice were placed on a rotarod (Bioseb) accelerating from 4 to 40 rpm over a span of 5 min. Mice were given 3 trials with a 30-sec inter-trial interval.

#### Immunostaining

Spinal cords from 2-wk-old mice were dissected and immersion-fixed for 1 h in 0.25% paraformaldehyde (PFA), then rinsed in phosphate buffered saline (PBS) and cryoprotected overnight in 20% sucrose at 4°C. Spinal cords were frozen in OCT medium (Tissue Tek) and 30 μm cryosections were collected from the L1-L2 and L4-L5 segments. Slices were rehydrated in PBS at room temperature for 15 min, permeated for 1 h in a blocking solution (BSA 1%, Normal Donkey Serum 3% in PBS) with 0.2% triton X-100, and for 12 h at 4 °C in a humid chamber with the primary polyclonal antibody anti-*Nav1.1* (1:400; #AB5204, Merck-Millipore from rabbit), or anti-*Nav1.6* (1:200; #ASC009, Alomone from rabbit) or anti-Choline Acetyltransferase (1/200; AB144P, Sigma-Aldrich) or anti-Ankyrin G (1:500; #SC-28561, Clinisciences from rabbit), or with the primary monoclonal anti-pan Na_v_ (1:1000, #S8809,Sigma from mouse) or anti-Ankyrin G (1:500; #MABN466, Merck-Millipore from mouse). Slices were then washed 3×5 min in PBS and incubated for 2 h with Alexa Fluor® 488-anti-mouse or 546-anti-rabbit or 546-anti-goat IgG secondary antibodies (1:800, 1:400, 1:400, #A-11017, #A-11071, A-11056, Lifetechnologies). After 3 washes of 5 min in PBS, they were mounted with a gelatinous aqueous medium. Sections were scanned using a laser scanning confocal microscope (Zeiss LSM700) in stacks of 0.3-1 μm-thick optical sections at × 63 and/or × 20 magnification respectively and processed with the Zen 12.0 software (Zeiss). Each optical section resulted from two scanning averages. Each figure corresponds to a projection image from a stack of optical sections. We used identical settings, finely tuned to avoid saturation, for the whole series.

#### Nav channels protein quantification

*<u>Membrane protein isolation and Western blots.</u>* Tissues were collected from spinal cord lumbar enlargements and frozen after removing the dorsal and ventral roots. For the membrane fraction, corresponding to the plasma membrane-enriched fraction, samples were homogenized in ice-cold lysis buffer (320 mM sucrose, 5 mM Tris-HCL pH 7.5, 10 µM iodoacetamide) supplemented with protease inhibitors (CompleteMini, Roche diagnostic Basel, Switzerland). Unsolubilized material was pelleted by centrifugation at 7,000 g for 5 min. The supernatant was subjected to an additional centrifugation step at 18000g for 70 min at 4 °C. Pellets were collected and homogenized in ice cold lysis buffer (1 % Igepal CA-630, Phosphate Buffer Saline 1X, 0.1% SDS, 10 µM iodoacetamide), supplemented with protease inhibitors (CompleteMini, Roche diagnostic). Protein concentrations were determined using a detergent-compatible protein assay (Bio-Rad). Equal protein amounts (30 µg) from samples were size fractionated by 4-15%

Mini-PROTEAN TGX stain-free gels (Bio-Rad), transferred to a nitrocellulose membrane and probed with the monoclonal anti-pan Na_v_ (1:500, #S8809, Sigma from mouse) at 4 °C overnight in Tris-buffered saline containing 5% fat-free milk powder. The blot was then incubated for 1 h at 22°C with a polyclonal horseradish peroxidase-conjugated anti-mouse IgG secondary antibody (1:40000; #31430, ThermoFisher). Signal intensities were measured with the image analysis software Quantity-One (BioRad).

#### Drug list and recording solutions

Normal aCSF was used in most cases for *in vitro* electrophysiological recordings (see above). Ca^2+^-free solution was made by removing Ca^2+^ chloride from the recording solution and replacing it with an equimolar concentration of magnesium chloride. Neurons were sometimes isolated from glutamatergic excitatory inputs with the kynurenic acid (1.5 mM). To isolate Na^+^ currents during voltage-clamp experiments we used a modified aCSF containing (in mM): 100 NaCl, 3 KCl, 1.25 NaH_2_PO_4_, 1.3 MgSO_4_, 3.6 MgCl_2_, 1.2 CaCl_2_, 25 NaHCO_3_, 40 D-glucose, 10 TEA-Cl and 0.1 CdCl_2_. All solutions were oxygenated with 95% O_2_/5% CO_2_. All salt compounds, tetraethylammonium (TEA; #T2265), N-methyl-DL-aspartic acid (NMA; #M2137), 5-hydroxytryptamine creatinine sulfate (5-HT; #S2805), were obtained from Sigma-Aldrich, 4,9-Anhydrotetrodotoxin (4,9-ahTTX; #6159) and ICA121431 (ICA; #5066/10) from Tocris Bioscience.

### Quantification and statistical analysis

#### Data analysis

*<u>Electrophysiological data</u>* were analyzed off-line with Clampfit 10.7 software (Molecular Devices). For whole-cell recordings, several basic criteria were set to ensure optimum quality of intracellular recordings. Only cells exhibiting a stable resting membrane potential, access resistance (no > 20% variation) and an action potential amplitude larger than 40 mV under normal aCSF were considered. Passive membrane properties of cells were measured by determining from the holding potential the largest voltage deflections induced by small current pulses that avoided activation of voltage-sensitive currents. We determined input resistance by the slope of linear fits to voltage responses evoked by small positive and negative current injections. Firing properties were measured from depolarizing current pulses of varying amplitudes. The rheobase was defined as the minimum step current intensity required to induce an action potential. We measured the action potential waveform properties of the first spike elicited by the rheobase sweep. The threshold for action potential was defined as the voltage at the point of deflection for dV/dt to be greater than zero. Peak spike amplitude was measured from the threshold potential, and spike duration was measured at half-amplitude. Action potential slopes (up and down) were measured as maximum slope during the action potential. The instantaneous discharge frequency was determined as the inverse of interspike interval. Firing frequency was calculated as the average action potential frequency over the entire duration of the pulse. All reported membrane potentials were corrected for the liquid junction potential. Bistable properties of motoneurons were investigated with a 2 s suprathreshold depolarizing current pulse elicited from a holding membrane potential at −60 mV. The cell was considered as bistable when (1) the pre-stimulus membrane potential stays below the spiking threshold (downstate), (2) the post-stimulus membrane potential stays depolarized (upstate), at peri-threshold for spike generation and (3) the membrane potential switches to downstate after a brief hyperpolarizing pulse. Bursting properties of interneurons were measured after smoothing to eliminate spikes but preserve the envelope of depolarization. Amplitude and duration of bursts were measured by a threshold function which determines the peak, the onset and the end of bursts. Burst frequency was calculated by dividing the number of bursts by the total duration of the recording. Voltage dependence and kinetics of *I*_NaP_ were analyzed from normalized voltage-ramp data by fitting them with Boltzmann functions. We defined the voltage-dependent activation threshold of the *I*_NaP_ as the membrane potential at which the slope of leak-subtracted current first becomes negative. We measured the magnitude of *I*_NaP_ as the peak of the leak-subtracted inward current during the ascending phase of the voltage command. For extracellular recordings, alternating activity between right/left L5 recordings was taken to be indicative of fictive locomotion. To characterize locomotor burst parameters, raw extracellular recordings from ventral roots were rectified, integrated and resampled at 50 Hz. Peak amplitude and duration of locomotor bursts were measured and the cycle period was calculated by measuring the time between the first two peaks of the auto-correlogram. *<u>Behavioral analyses:</u>* For walking, bottom view recordings were processed by the CatWalk XT software (Noldus Information Technology, Netherlands) to measure a broad number of spatial and temporal gait parameters in several categories. These include (1) the paw print area that refers to the total area of the glass in contact with the paw, measured when the area of contact with the glass is maximal; (2) dynamic parameters related to individual paw prints, such as duration of the step cycle; (3) parameters related to the position of paw prints with respect to each other, for example the hindlimb base of support (the width between the paw pairs); (4) parameters related to time-based relationships between paw pairs. These parameters were calculated for each run and for each paw. To estimate five hindlimb joints positions (toe, ankle, knee, hip and iliac crest) on lateral view videos of freely walking mice, we trained a model using the DeepLabcut toolbox. A total of 320 image frames (16 videos, 20 frames/video) were used to label and train the model. A pre-trained network with 50 layers (ResNet50) was selected and trained for 200 000 iterations. Parameters describing gait timing and joint kinematics were extracted for each gait cycle using a custom-written Python script. Gait cycles were defined as the time interval between two successive paw contacts of one limb. Individual steps were identified within the run by the acceleration of the paws to identify the “stance” and “swing” phases. We measured the total angular excursion for the hip, knee and ankle joints across the cycle for the hindlimbs. In the rotarod test the average latency of the subject to fall was recorded (in seconds). All behavioral tests were carried out with the experimenter blind to the experimental group.

*<u>Immunohistochemistry analysis:</u>* Quantification of staining intensities were performed with the Zen 12.0 software (Zeiss). Measurements were performed on the axon initial segment (AIS) from interneurons located in the ventromedial part of upper lumbar segments (L_1_-L_2_) near the central canal and from motoneurons identified as the biggest cells located in the ventral horn (L_4_-L_5_). The AIS was identified as linear structures labeled by pan Na_v_-specific antibodies, and for which the beginning and the end of the structure could be clearly determined. Immunofluorescence intensity profiles were obtained from a line traced along the AIS. For each staining, the background intensity was first subtracted, and immunofluorescence values along each AIS were then normalized to the maximum intensity. The Pearson correlation coefficient was used to quantify the degree of colocalization between fluorophores. The measurements were repeated with similar numbers of motoneurons per animal.

#### Statistics

No statistical method was used to predetermine sample size. Group measurements were expressed as means ± s.e.m. When two groups (control vs transgenic mice) were compared, we used the unpaired Mann–Whitney test or the Unpaired *t*-test. Fisher test was used to compare the percentages of bistability. When two conditions (control vs drugs) were compared, we used the Wilcoxon matched pairs test. We also used the one-way or two-way ANOVA tests for multiple comparisons. For all statistical analyses, the data met the assumptions of the test and the variance between the statistically compared groups was similar. The level of significance was set at p < 0.05. Statistical analyses were performed using Graphpad Prism 7 software and are indicated in Figure legends.

## RESOURCE TABLE

**Key Resources Table.**
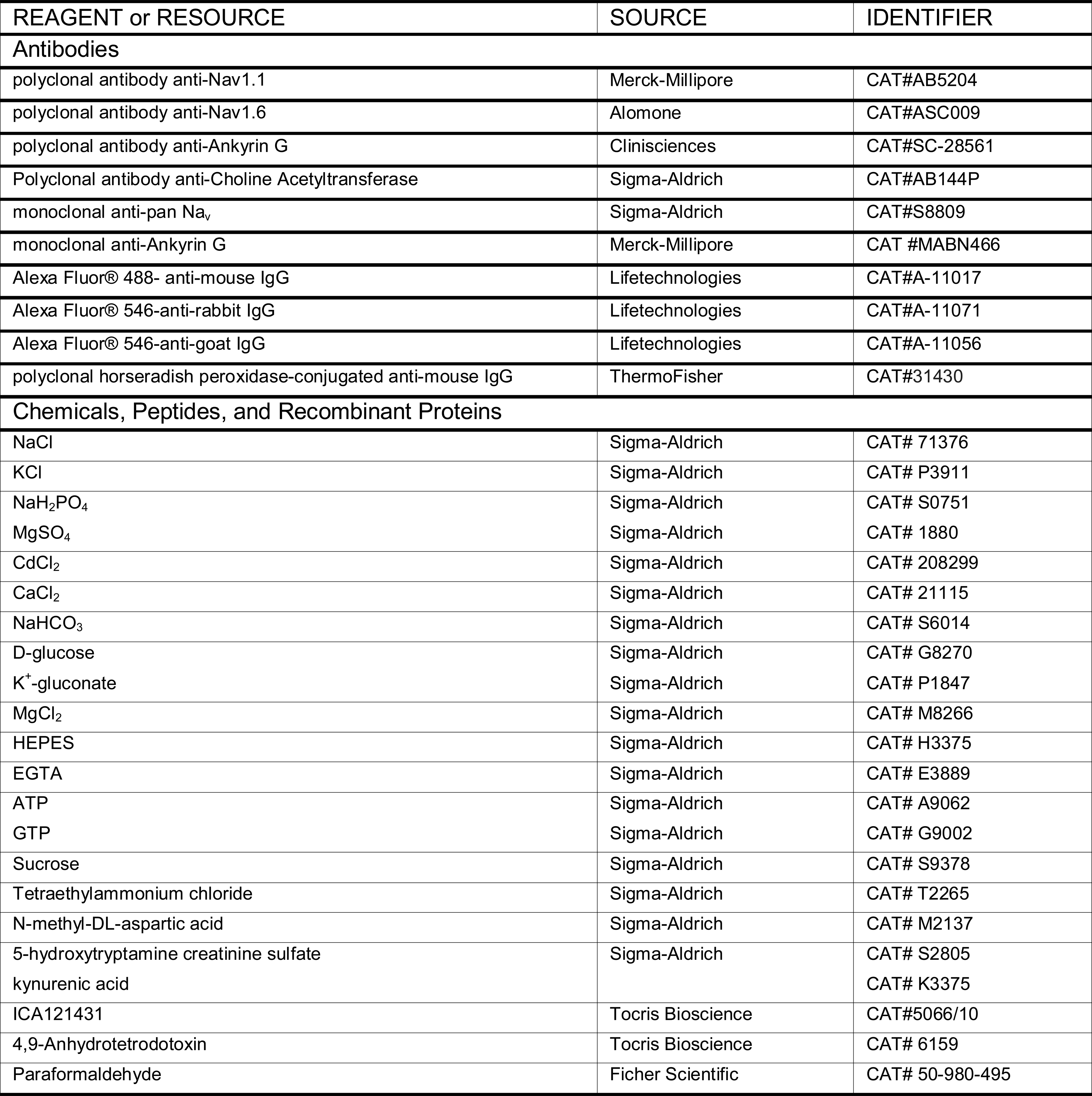
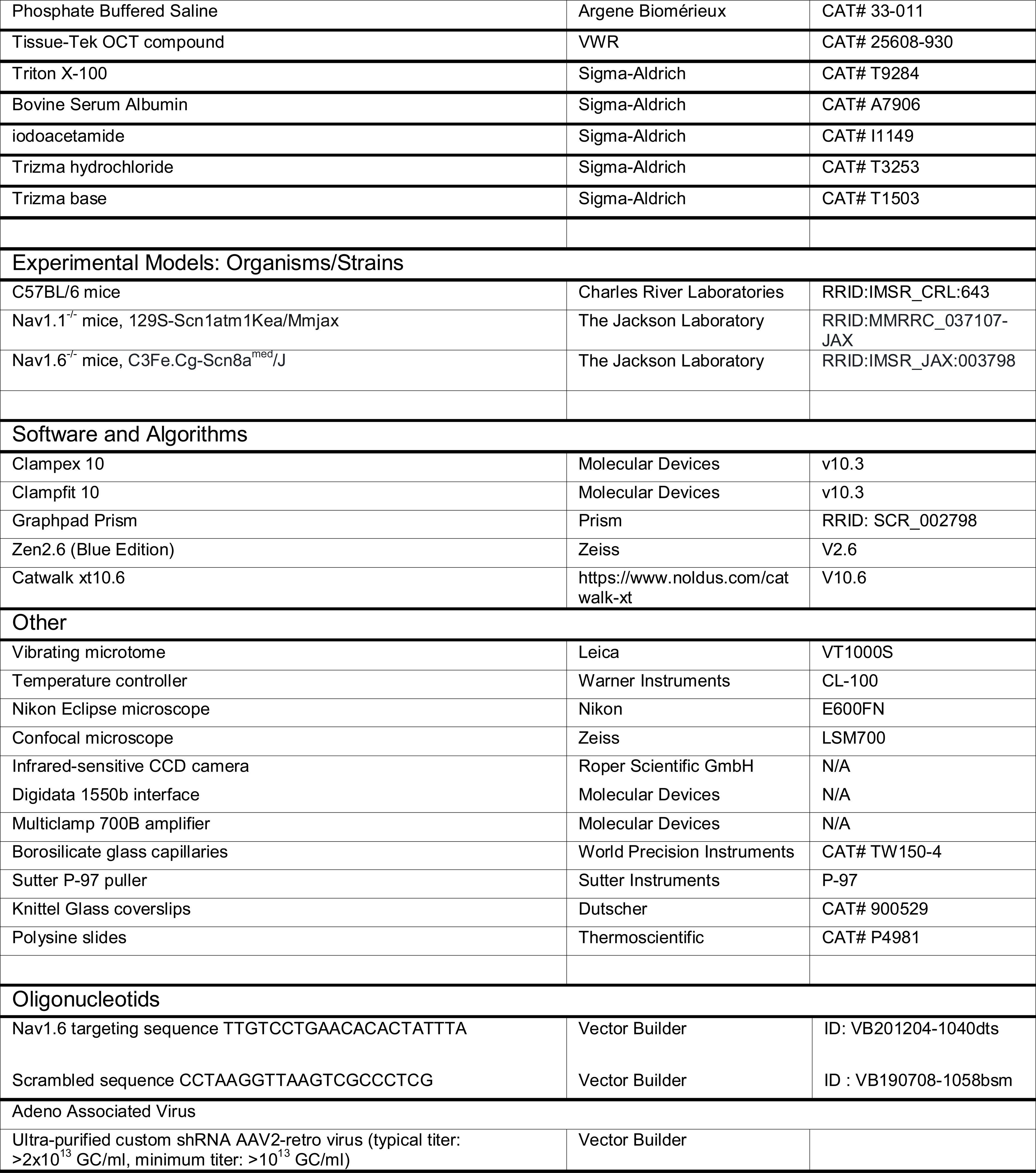

