## Supplemental information for "Persistent Nav1.6 current drives spinal locomotor functions through nonlinear dynamics"

**Supplementary information**

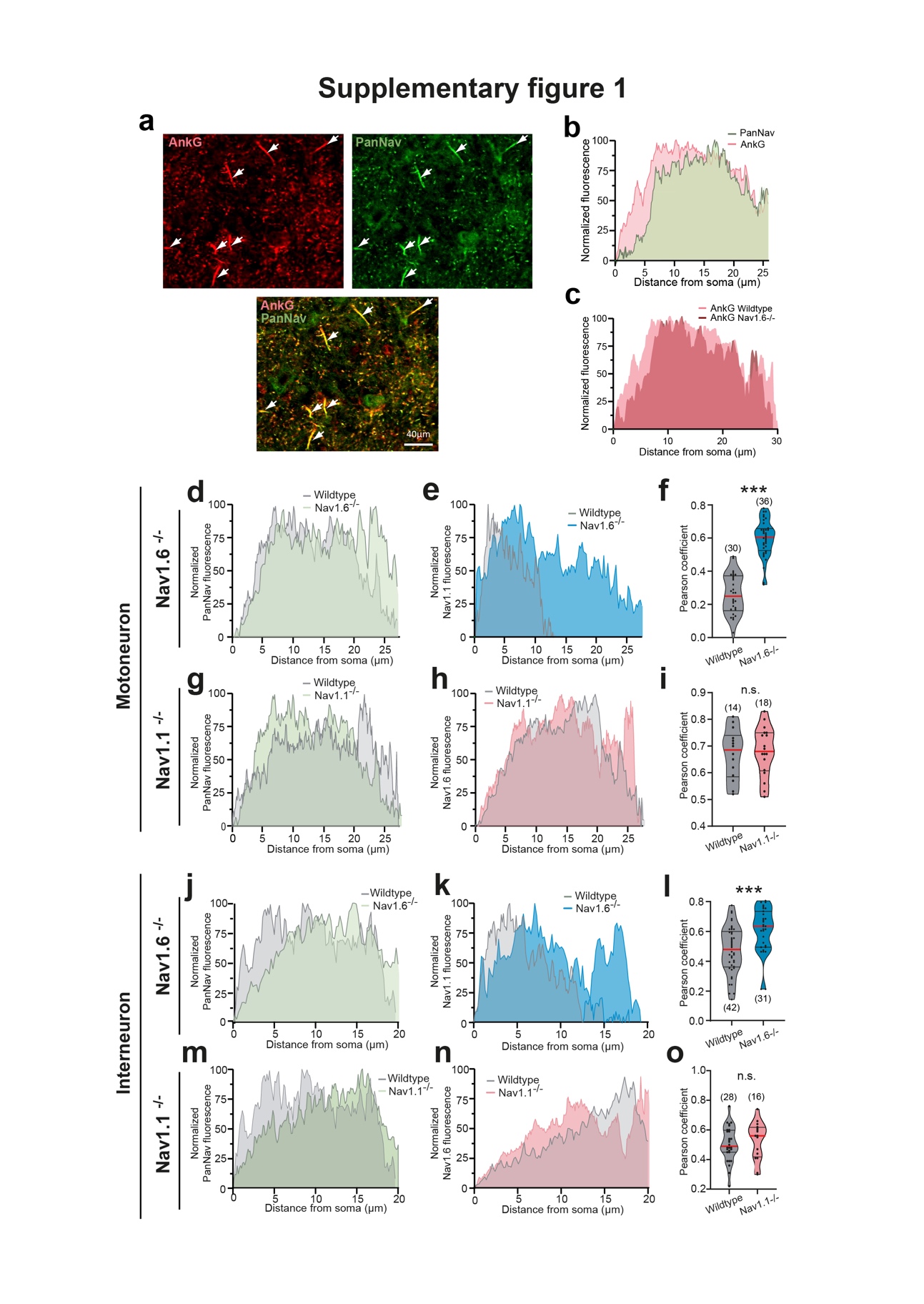

**Supplementary Fig. 1 (related to Fig.1): Distribution of Nav channels along the axon initial segments (AISs) of spinal neurons. a** Confocal images of the ventral horn double-labelled with anti-ankyrin-G (top-left, red) and PanNav (top-right, green) antibodies; Merge image (bottom panel); Scale bar represent 40 µm. Arrowheads point to axon initial segments (AISs) from presumptive motoneurons. **b** Mean fluorescence intensity profile for the PanNav (green), along the ankyrin-G-labelled AIS (red) of motoneurons. **c** Mean fluorescence intensity profile for the ankyrin-G along the AIS of motoneurons from wildtype (pink) and *Nav1.6*^-/-^ (red) mice. **d,e,g,h,j,k,m,n** Mean fluorescence intensity profile for the PanNav (**d,g,j,m**), *Nav1.1* (**e,k**) and *Nav1.6* (**h,n**) immunostaining along the PanNav-labelled AIS of both motoneurons (**d,e,g,h**) and interneurons (**j,k,m,n**) from *Nav1.1*^-/-^ (**g,h,m,n**) and *Nav1.6*^-/-^ (**d,e,j,k**) mice. For each antibody, the immunofluorescence was normalized to its maximum intensity. **f,i,l,o** Violin plots of the Pearson’s coefficient between PanNav and Na1.1 (**f,l**) or between PanNav and *Nav1.6* (**i,o**) in motoneurons (**f,i**) and interneurons (**l,o**) from *Nav1.1*^-/-^ (**i,o**) and *Nav1.6*^-/-^ (**f,l**) mice. Numbers in brackets in **f,i,l,o** indicate the numbers of cells. Each dot represents an individual neuron. n.s., no significance; ****P* < 0.001 (Unpaired t-test for **f,i,l,o**). For detailed *P* values see Source Data. Source data are provided as a Source data file.

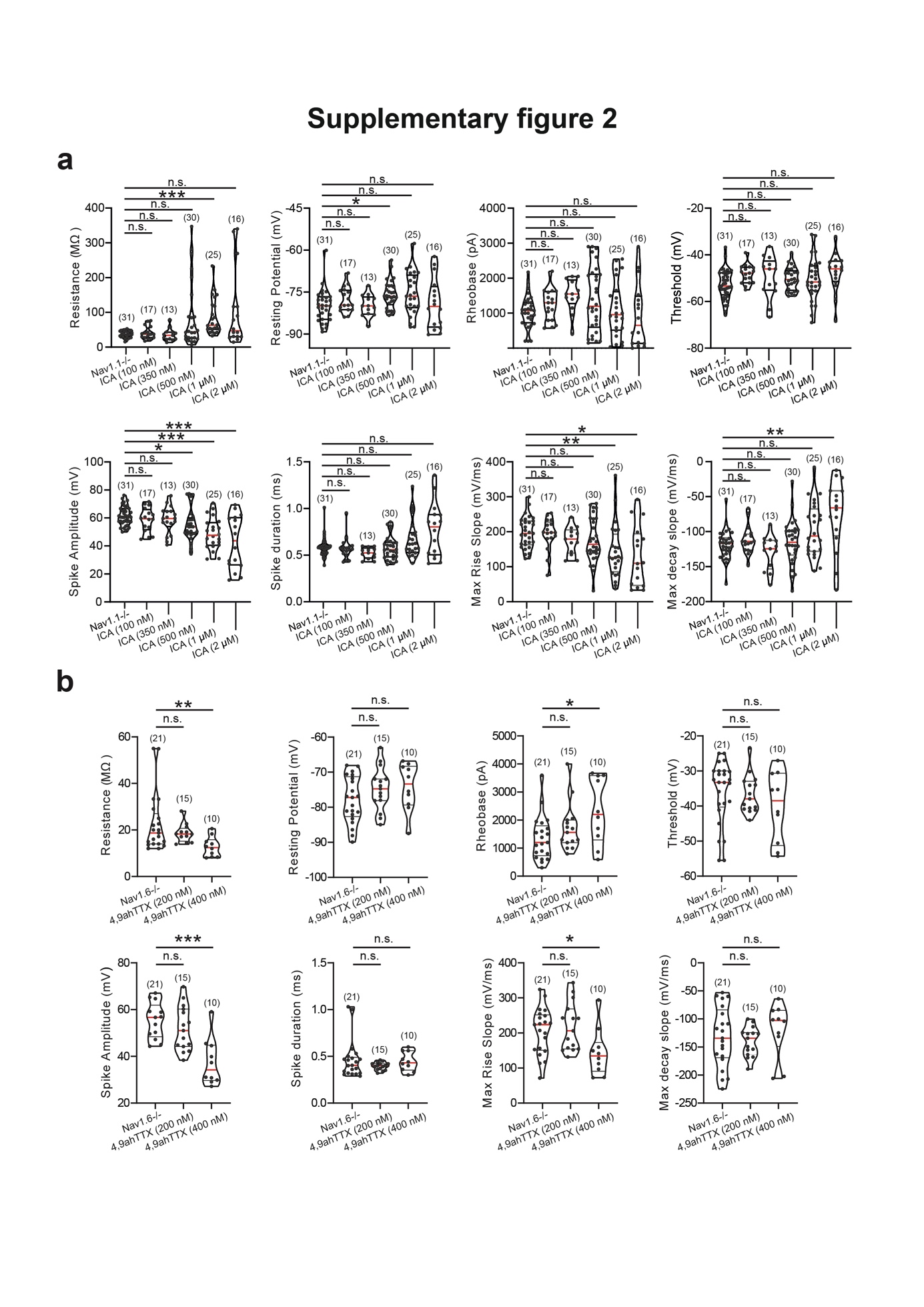

**Supplementary Fig. 2 (related to Fig. 2): Determination of the optimal concentration of ICA-121431 and 4,9-anhydrotetrodotoxin to inhibit *Nav1.1* and *Nav1.6* channels, respectively. a,b** Violin plots quantifying the effects of ICA-121431 (**a,** ICA, *n* = 9 mice) and 4,9-anhydrotetrodotoxin (**b,** 4,9 ahTTX, *n* = 5 mice) across different concentrations on passive and active membrane properties of *Nav1.1*^-/-^ and *Nav1.6*^-/-^ motoneurons, respectively. Numbers in brackets indicate the numbers of motoneurons. Each dot represents an individual motoneuron. n.s., no significance; **P* < 0.05; ***P* < 0.01; ****P* < 0.001 (one-way ANOVA with multiple comparisons for **a** and **b**). For detailed *P* values, see Source data. Source data are provided as a Source data file.

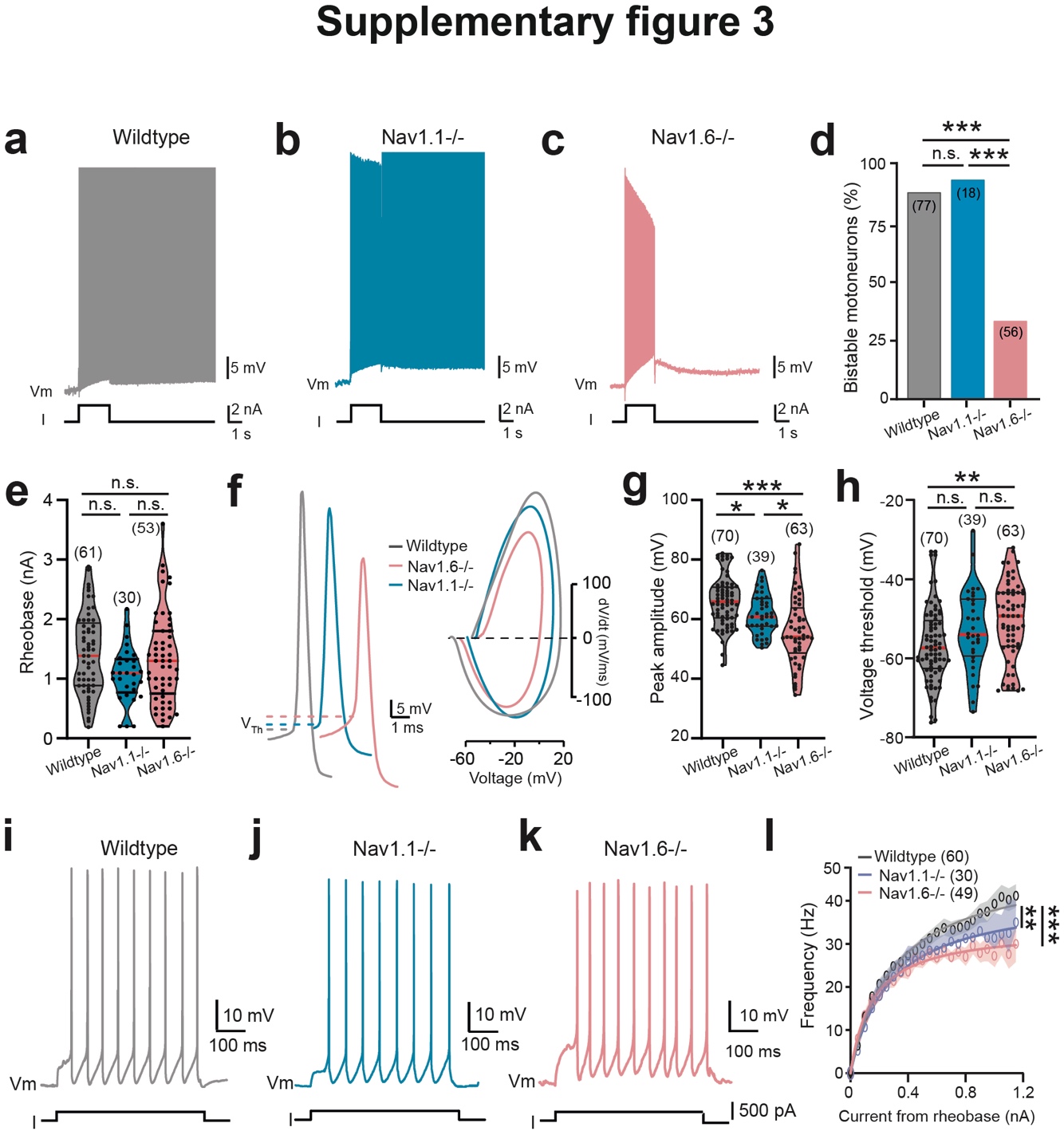

**Supplementary Fig. 3 (related to Fig. 2): Altered bistability of lumbar motoneurons in *Nav1.6*^-/-^ mice. a-c,f,i-k** Voltage traces from motoneurons in response to suprathreshold (**a-c**), near-threshold (**f**) or incrementing (**i-k**) depolarizing pulses recorded from wildtype (**a**,**f,i,** grey, *n* = 6 mice) *Nav1.1*^-/-^ (**b,f,j**, blue, *n* = 5 mice) or *Nav1.6*^-/-^ (**c,f,k**, pink, *n* = 7 mice) mice. **d** Quantification of the proportion of bistable motoneurons. **e,g,h** Violin plots of the rheobase (**e**), peak amplitude (**g**) and threshold (**h**) of the action potential. **f** Representative individual action potentials (left) recorded in motoneurons from wildtype (grey), *Nav1.1*^-/-^ (blue) or *Nav1.6*^-/-^ (pink) mice. with their phase plots (right) generated from the first derivative (dV/dt; y-axis) versus membrane potential (mV; x-axis). Dashed lines indicate the spiking threshold (V_Th_). **l** Firing frequency as a function of the amplitude of the current pulse. Continuous lines represent best fit functions for experimental data with 95% confidence interval. Numbers in brackets in **d,e,g,h,l** indicate the numbers of motoneurons. Each dot represents an individual cell. n.s., no significance; ***P* < 0.01; ****P* < 0.001 (two-tailed Fisher test for **d**; one-way ANOVA with multiple comparisons for **e,g,h**; comparison of the fits for **l**). For detailed *P* values, see Source data. Source data are provided as a Source data file.

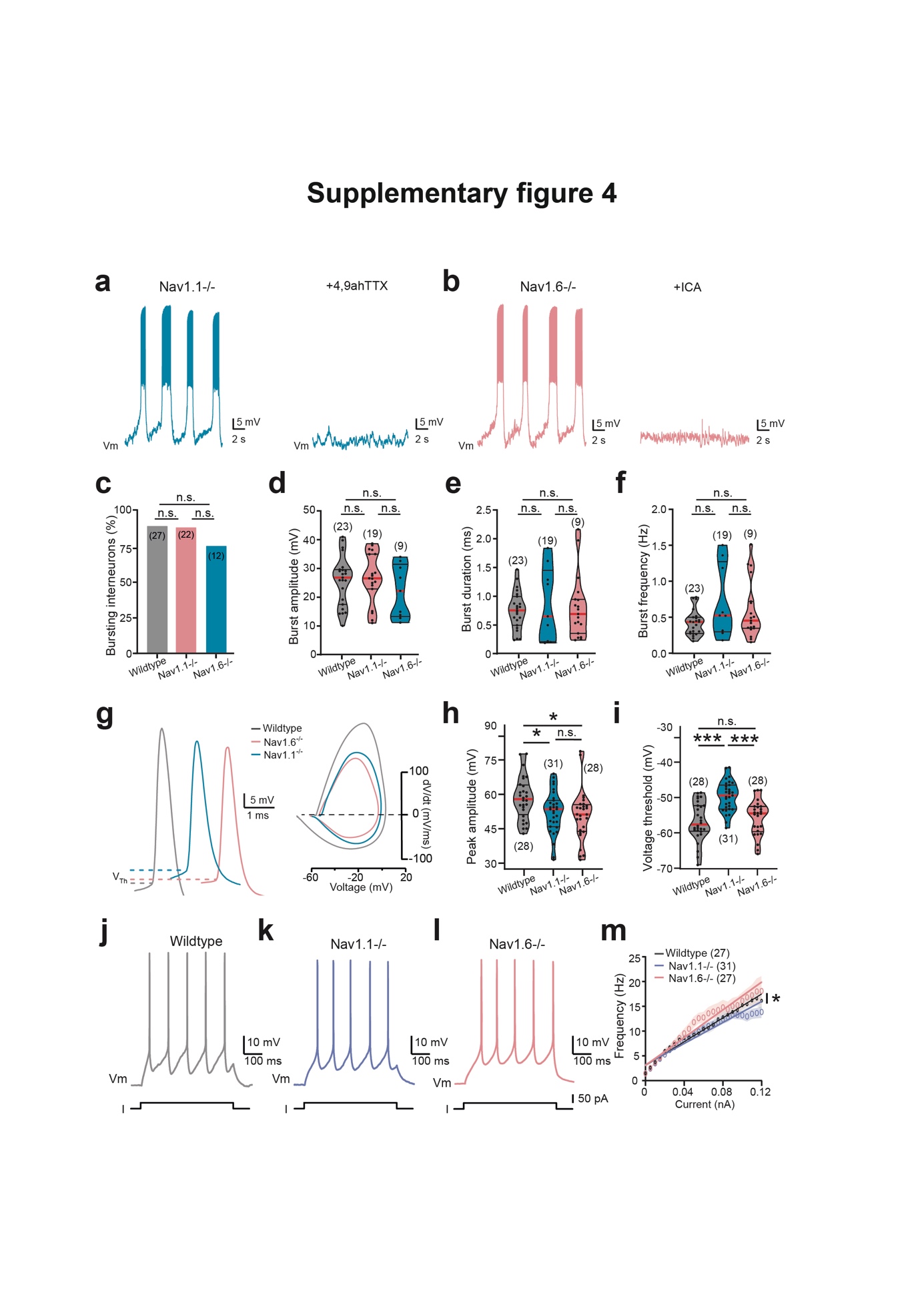

**Supplementary Fig. 4 (related to Fig. 3): *I*_NaP_-dependent bursting properties in mutant mice.** **a,b** [Ca^2+^]_o_-free-saline-induced bursting activity recorded from ventromedial interneurons of the locomotor CPG region (L_1_-L_2_) in *Nav1.1*^-/-^ mice (**a**) before (left) and during (right) the bath application of 4,9-anhydrotetrodotoxin (4,9 ahTTX, 200 nM, *n* = 2 mice) or in *Nav1.6*^-/-^ mice (**b**) before (left) and during (right) the bath application of ICA-121431 (ICA, 350 nM, *n* = 2 mice). **c** Quantification of the proportion of bursting cells. **d-f** Quantification of burst parameters. **g** Representative individual action potentials (left) recorded in interneurons from wildtype (grey), *Nav1.1*^-/-^ (blue) or *Nav1.6*^-/-^ (pink) mice with their phase plots (right) generated from the first derivative (dV/dt; y-axis) versus membrane potential (mV; x-axis). Dashed lines indicate the spiking threshold (V_Th_). **h,i** Violin plots of the peak amplitude (**h**) and threshold (**i**) of the action potential. **j-l** Voltage traces from interneurons in response to incrementing depolarizing pulses recorded from wildtype (**j,** *n* = 7 mice), *Nav1.1*^-/-^ (**k,** *n* = 3 mice) or *Nav1.6*^-/-^ (**l,** *n* = 7 mice) mice. **m** Firing frequency as a function of the amplitude of the current pulse. Continuous lines represent best fit functions for experimental data with 95% confidence interval. Numbers in brackets in **c-f,h,i,m** indicate the number of interneurons. Each dot represents an individual cell. n.s., no significance; **P* < 0.05; ****P* < 0.001 (two-tailed Fisher test for **c**; one-way ANOVA with multiple comparisons for **d-f,h,i**; comparison of the fits for **m**). For detailed *P* values, see Source data. Source data are provided as a Source data file.

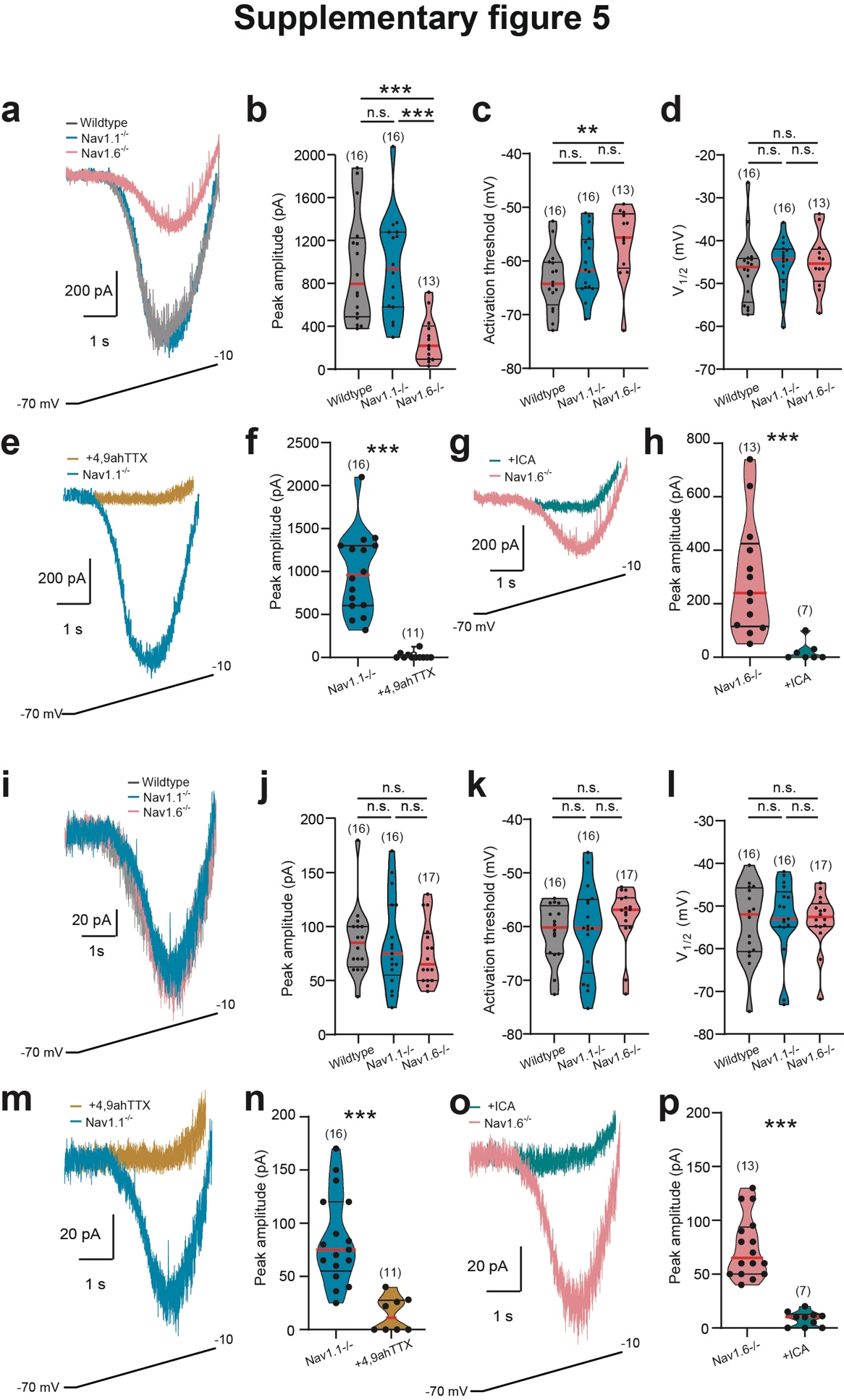

**Supplementary Fig. 5 (related to Fig. 4): Motoneurons lacking the *Nav1.6* subunits display reduced *I*_NaP_**. **a,i** Leak-subtracted *I*_NaP_ recorded in motoneurons (**a**) and interneurons of the CPG region (**i**) from wildtype (grey, *n* = 3 mice), *Nav1.1*^-/-^ (blue, *n* = 3 mice) and *Nav1.6*^-/-^ (pink, *n* = 2 mice) mice in response to a slow ramping depolarization. **e,m** Leak-subtracted *I*_NaP_ recorded in motoneurons (**e**) and interneurons of the CPG region (**m**) from *Nav1.1*^-/-^ mice before and during the bath application of 4,9-anhydrotetrodotoxin (4,9 ahTTX, 200 nM). **g,o** Leak-subtracted *I*_NaP_ recorded in motoneurons (**g**) and interneurons of the CPG region (**o**) from *Nav1.6*^-/-^ mice before and during the bath application of ICA-121431 (ICA, 350 nM). **b-d,f,h,j-l,n,p** Quantification of biophysical properties of *I*_NaP_. Numbers in brackets indicate the number of neurons. Each dot represents an individual neuron. n.s., no significance; ***P* < 0.01; ****P* < 0.001 (two-tailed Mann-Whitney test for **f,h,n,p**; one-way ANOVA with multiple comparisons for **b-d,j-l**). For detailed *P* values, see Source data. Source data are provided as a Source data file.

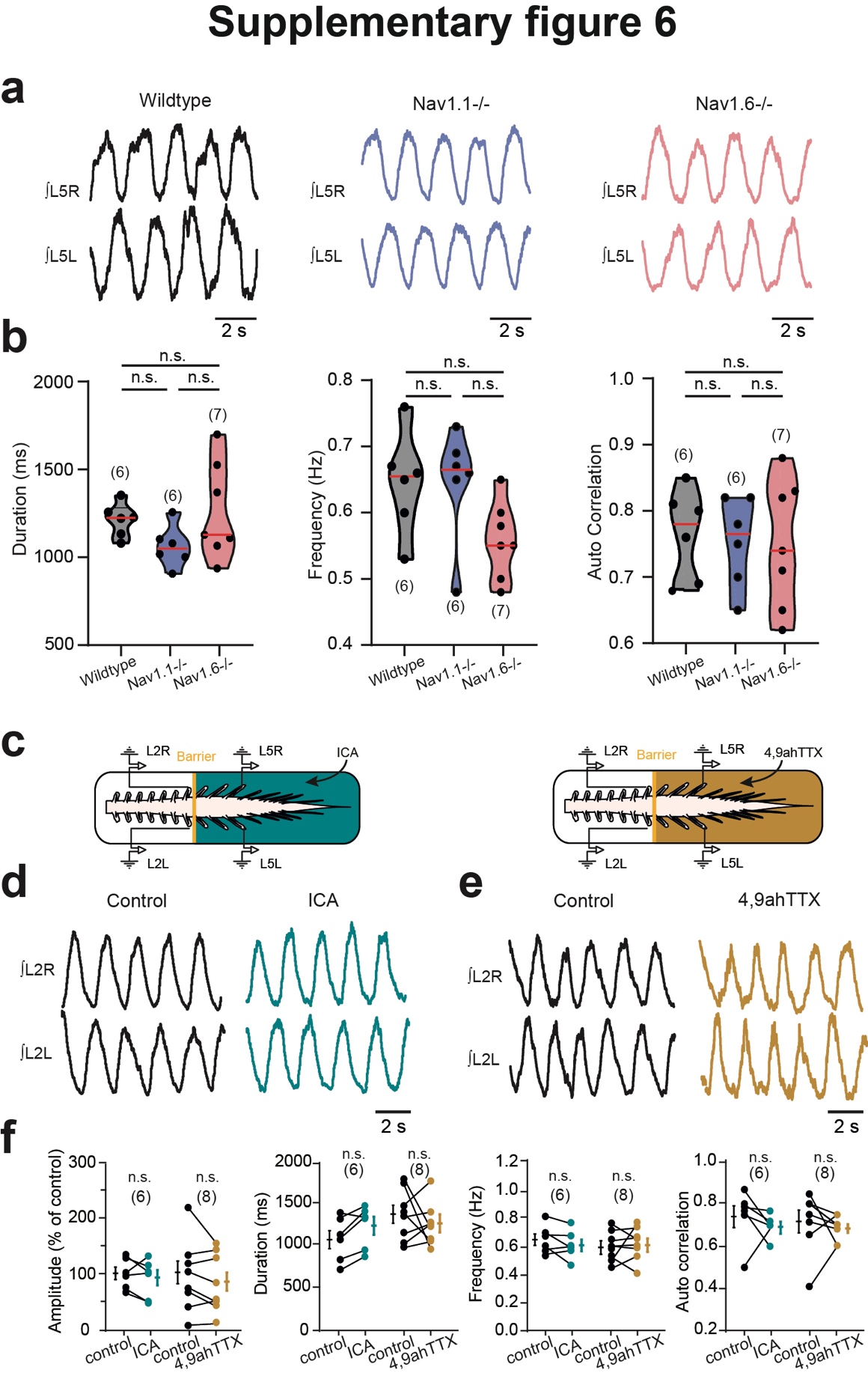

**Supplementary Fig. 6 (related to Fig. 5): Spinal cords isolated from *Nav1.1*^-/-^ and *Nav1.6*^-/-^ mutant mice display normal fiction locomotion.** **a** L5 Ventral-root recordings of NMA/5-HT-induced fictive locomotor activity recorded in spinal cords isolated from wildtype (left, black, *n* = 6 mice), *Nav1.1*^-/-^ (middle, blue, n = 6 mice) or *Nav1.6*^-/-^ (right, pink, n = 7 mice) mice. **b,f** Quantification of locomotor burst parameters. **c** Schematic representation of the whole-mount spinal cord with the recording glass electrodes from homosegmental L_2_-L_5_ (R/L). The yellow solid line represents the Vaseline barrier. Built at L_2_/L_3_ level the Vaseline barrier allows the selective application of drugs over the rostral (above L_3_ segment) or the caudal lumbar segments (below L_3_ segment). **d,e** L2 Ventral-root recordings of NMA/5-HT-induced fictive locomotor activity recorded in spinal cords isolated from wildtype mice before and after adding ICA-121431 (**d**, ICA, 350 nM, *n* = 6 mice) or 4,9-anhydrotetrodotoxin (**e**, 4,9 ahTTX, 200 nM, *n* = 8 mice) to caudal lumbar segments. Numbers in brackets in **b,f** indicate the number of spinal cords. Each dot represents an individual spinal cord. n.s., no significance (one-way ANOVA with multiple comparisons for **b**; two-tailed Wilcoxon paired test for **f**). Mean ± SEM. For detailed *P* values, see Source data. Source data are provided as a Source data file.

**Supplementary Movie 1:** Representative raw video from 4 wk-old control-shRNA mice freely walking through a corridor.

**Supplementary Movie 2:** Representative raw video from 4 wk-old *Nav1.6*-shRNA mice freely walking through a corridor.
